# Somatic cancer driver mutations are enriched and associated with inflammatory states in Alzheimer’s disease microglia

**DOI:** 10.1101/2024.01.03.574078

**Authors:** August Yue Huang, Zinan Zhou, Maya Talukdar, Michael B. Miller, Brian Chhouk, Liz Enyenihi, Ila Rosen, Edward Stronge, Boxun Zhao, Dachan Kim, Jaejoon Choi, Sattar Khoshkhoo, Junho Kim, Javier Ganz, Kyle Travaglini, Mariano Gabitto, Rebecca Hodge, Eitan Kaplan, Ed Lein, Philip L. De Jager, David A. Bennett, Eunjung Alice Lee, Christopher A. Walsh

## Abstract

Alzheimer’s disease (AD) is an age-associated neurodegenerative disorder characterized by progressive neuronal loss and pathological accumulation of the misfolded proteins amyloid-β and tau^1,2^. Neuroinflammation mediated by microglia and brain-resident macrophages plays a crucial role in AD pathogenesis^1–5^, though the mechanisms by which age, genes, and other risk factors interact remain largely unknown. Somatic mutations accumulate with age and lead to clonal expansion of many cell types, contributing to cancer and many non-cancer diseases^6,7^. Here we studied somatic mutation in normal aged and AD brains by three orthogonal methods and in three independent AD cohorts. Analysis of bulk RNA sequencing data from 866 samples from different brain regions revealed significantly higher (∼two-fold) overall burdens of somatic single-nucleotide variants (sSNVs) in AD brains compared to age-matched controls. Molecular-barcoded deep (>1000X) gene panel sequencing of 311 prefrontal cortex samples showed enrichment of sSNVs and somatic insertions and deletions (sIndels) in cancer driver genes in AD brain compared to control, with recurrent, and often multiple, mutations in genes implicated in clonal hematopoiesis (CH)^8,9^. Pathogenic sSNVs were enriched in CSF1R+ microglia of AD brains, and the high proportion of microglia (up to 40%) carrying some sSNVs in cancer driver genes suggests mutation-driven microglial clonal expansion (MiCE). Analysis of single-nucleus RNA sequencing (snRNAseq) from temporal neocortex of 62 additional AD cases and controls exhibited nominally increased mosaic chromosomal alterations (mCAs) associated with CH^10,11^. Microglia carrying mCA showed upregulated pro-inflammatory genes, resembling the transcriptomic features of disease-associated microglia (DAM) in AD. Our results suggest that somatic driver mutations in microglia are common with normal aging but further enriched in AD brain, driving MiCE with inflammatory and DAM signatures. Our findings provide the first insights into microglial clonal dynamics in AD and identify potential new approaches to AD diagnosis and therapy.

## Main-text

The importance of microglia in AD pathogenesis has been demonstrated by large-scale genetic association studies which have identified risk variants in a growing list of microglia-related genes^12–15^. As the primary immune cells in the central nervous system (CNS), microglia play critical roles in brain development, injury response, and pathogen defense^16^, modulating cellular responses involved in aging and neurodegeneration as well^3–5^. Once abnormally reactive in AD, microglia can promote synaptic and neuronal loss and exacerbate tau proteinopathy^17,18^. Recent single-cell transcriptomic studies have depicted specific populations of microglia enriched in AD brains of mouse models and human patients, termed disease-associated microglia (DAM)^19^. DAM feature reduced expression of homeostatic genes but elevated expression of genes involved in immune response and phagocytosis^3,20^, though whether DAM are beneficial or detrimental to AD remains unsettled^21^.

Somatic mutations accumulate in all cell types that have been studied, both during normal development and during aging^22–24^. Clonal expansion, driven by somatic mutations in genes regulating cell proliferation, is considered the major cause of cancer^6^, but has also been recently reported in various non-cancer cell types^7^ often in the absence of visible pathology. Clonal expansion of mutant blood cells, called clonal hematopoiesis (CH), increases in prevalence with age and is associated with increased risk of hematologic malignancies and cardiovascular disease^8,9^, likely through inflammatory effects of mutant cells on neighboring nonmutant cells^25^. A somatic V600E mutation in *BRAF*, a common cancer-driver mutation, in the microglial lineage has also been causally implicated in degeneration of neurons secondary to mutant microglial activation in both mouse models and humans^26^. Although gene panel sequencing of 20 AD brains^27^ and whole exome sequencing of DNA from micro-dissected neuronal nuclei of 52 AD brains^28^ found no consistent excess of clonal somatic mutations in AD, these studies were extremely limited in their ability to detect clonal somatic mutations by small sample sizes, the examination of neuronal DNA only, and low sequence coverage.

Here we tested whether brain clonal somatic mutation is associated with AD by three prospective and orthogonal approaches in >600 AD samples and >500 control brains of three AD cohorts (Fig. 1a-c), and we found consistent increases in overall clonal somatic mutations in AD compared to control, as well as function-specific enrichment in genes previously implicated in CH and other pre-cancerous conditions. These somatic mutations were enriched in microglia compared to other brain cell types, and microglia harboring these mutations exhibited a pro-inflammatory transcriptional signature that has previously been associated with neurodegeneration.

**Fig. 1.**
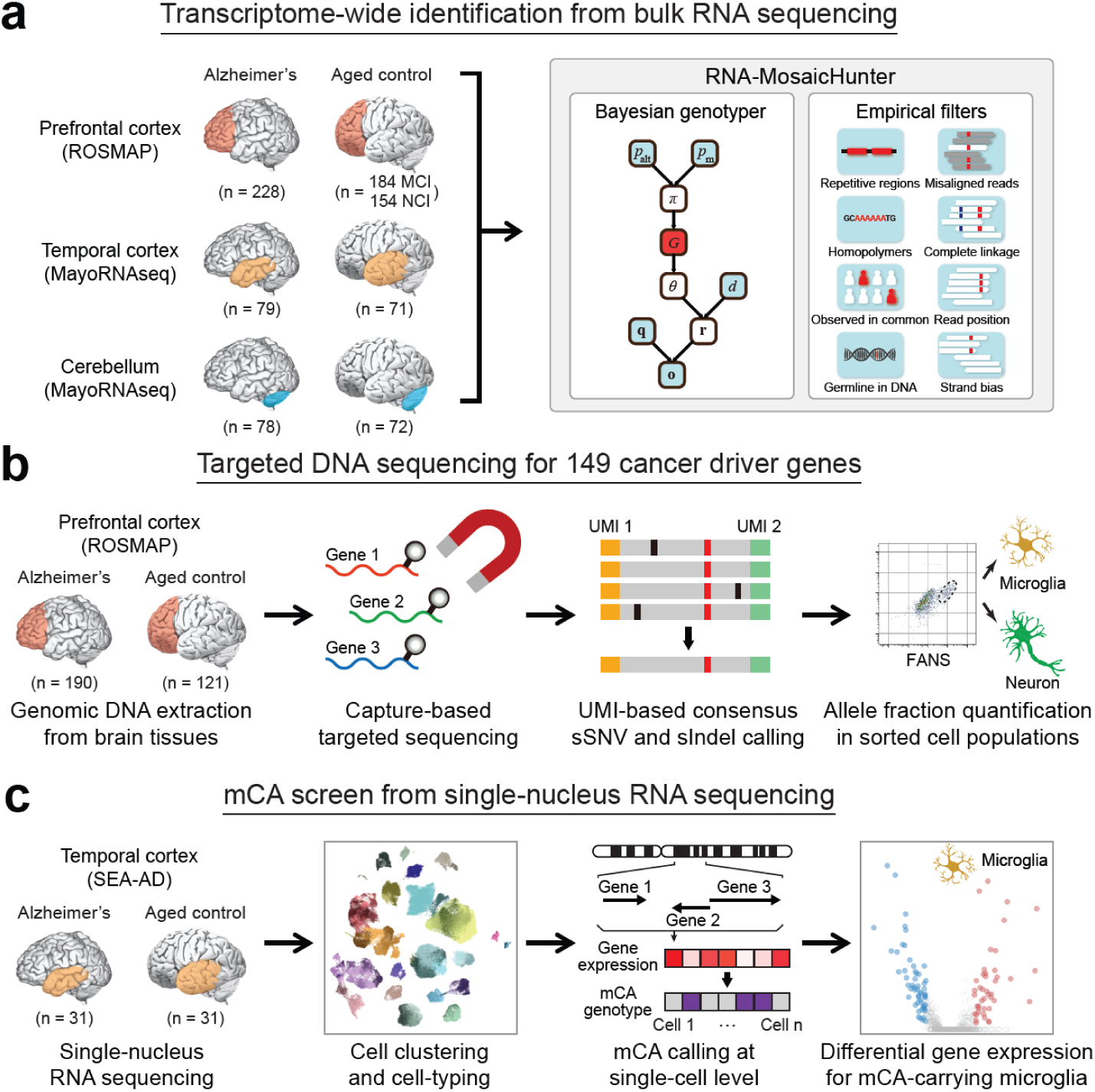
Overview of the experimental and analysis strategies. **a**, Transcriptome-wide screen of sSNVs among 886 bulk RNA-seq data sets of AD and control brain samples. Somatic mutations were called by RNA-MosaicHunter. MCI, mild cognitive impairment; NCI, no cognitive impairment. **b**, Profiling sSNVs and sIndels in 311 AD and control PFC samples using deep molecular barcode sequencing with a panel of 149 cancer driver genes. Mutation candidates were validated by amplicon sequencing and their mutant allele fractions were measured in different FANS-sorted nuclei populations. **c**, Identification and transcriptomic profiling of microglia in AD and control brain single-nucleus RNA-seq samples carrying mCA.

### Identifying somatic mutations from bulk RNA sequencing

We first developed RNA-MosaicHunter, a method to identify somatic mutations in coding regions of expressed genes, and applied it to 866 bulk RNA sequencing (RNA-seq) data sets of various brain regions including prefrontal cortex (PFC), temporal cortex, and cerebellum (Fig. 1a). The RNA-seq datasets were obtained from two independent harmonized cohorts of aging and dementia, the Rush Religious Orders Study/Memory and Aging Project (ROSMAP)^29^ and a collection of brains under the Mayo Clinic Alzheimer’s Disease Genetics Studies (MayoRNAseq)^30^, in which the clinical consensus diagnosis of cognitive status was given by expert neurologists based on detailed cognitive and neuropathologic phenotyping.

RNA-MosaicHunter, an extensive modification of MosaicHunter^31^, developed for sSNV calling in various types of DNA sequencing (DNA-seq) data, first calculates the likelihood of somatic mutation for each genomic position using a Bayesian graphical model, which distinguishes true mutations from random sequencing errors by considering base quality metrics for covered reads (Fig. 1a). RNA-MosaicHunter also incorporates a series of empirical filters to remove artifacts due to systematic base-calling and alignment errors in RNA-seq. Germline variants were removed by comparing against matched whole-genome or whole-exome sequencing data of the same individual. Considering the widespread adenosine-to-inosine (A-to-I) RNA editing sites across the genome^32^, where inosine will be recognized as guanine (G) and therefore indistinguishable from A-to-G sSNVs in RNA-seq data, we only considered non-A-to-G sites as sSNV candidates.

We benchmarked RNA-MosaicHunter using 19 esophageal carcinoma samples obtained from The Cancer Genome Atlas (TCGA) Research Network^33^. RNA-MosaicHunter identified 613 non-A-to-G sSNVs from the RNA-seq data, and 513 of them were supported by MuTect^34^ calls in matched whole-exome sequencing data, confirming the accuracy of RNA-MosaicHunter (Fig. 2a). In addition, 65 of 100 sSNVs that were detected by RNA-MosaicHunter but not MuTect showed mutant-supporting reads with >2% mutant allele fraction (MAF) in the DNA-seq data, suggesting that they were true somatic mutations omitted by MuTect (Fig. 2a). Among 851 MuTect-called exonic mutations with sufficient RNA-seq read coverage, RNA-MosaicHunter successfully recaptured 499 of them (Fig. 2b). In summary, RNA-MosaicHunter achieved 59% sensitivity and 94% precision to identify non-A-to-G sSNVs from the tumor RNA-seq data (Fig. 2b); the sSNVs missed by RNA-MosaicHunter generally had poor coverage or low MAF in RNA-seq data, likely due to their low expression level or allele-specific expression^35^ in the tumor samples.

**Fig. 2.**
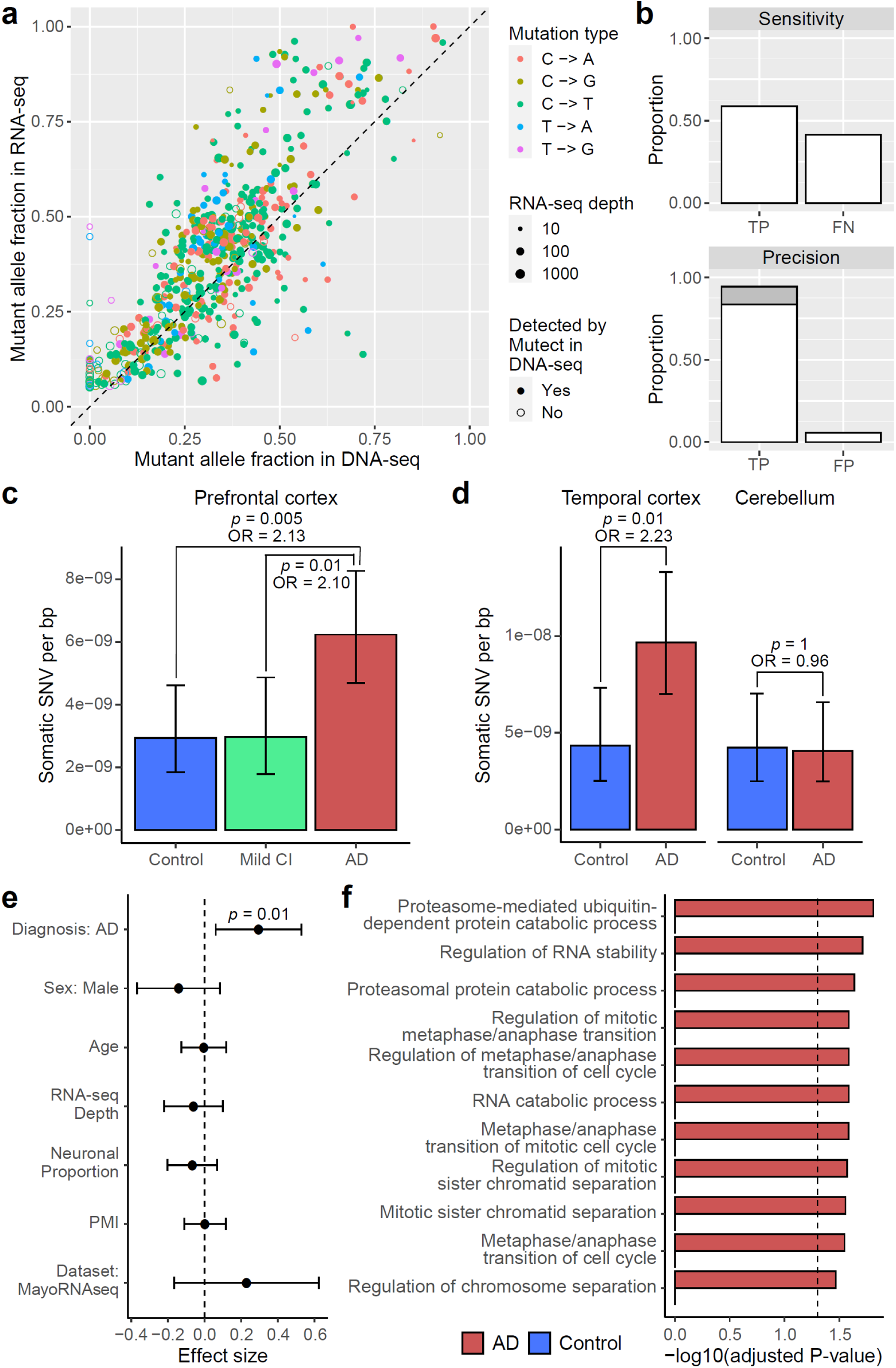
RNA-MosaicHunter reveals elevated burden of somatic mutations in the cerebral cortex of AD patients. **a**-**b**, Benchmarking the performance of RNA-MosaicHunter using the TCGA cancer data. 513 of 613 sSNVs identified by RNA-MosaicHunter were confirmed by MuTect in the matched DNA-seq data (filled circle in **a**). RNA-MosaicHunter recaptured 65 sSNVs that are present in DNA-seq but missed by MuTect (open circle in **a**; grey bar in **b**). TP, true positive; FN, false negative; FP, false positive. **c-d**, Greater mutation burden in cerebral cortex samples of AD patients when compared to matched controls. A significant two-fold increase of sSNV density in AD prefrontal cortex and temporal cortex was consistently found in both ROSMAP (**c**) and MayoRNAseq (**d**) cohorts. The burden increase was not observed in the AD cerebellum. CI, cognitive impairment. **e**, Linear regression modeling confirms that the sSNV increase in AD brains remains significant after controlling for potential covariates. PMI, post-mortem interval. **f**, Gene Ontology terms enriched for AD sSNVs. Genes regulating cell cycle and proliferation are specifically enriched for AD but not control sSNVs. **c**-**e**, Error bar, 95% CI.

### Higher burden of somatic mutation in AD cortex

RNA-MosaicHunter revealed two-fold increases in clonal somatic mutations compared to matched controls in two different AD cohorts. In PFC RNA-seq data of 228 persons with AD and 338 non-AD controls (Extended Data Fig. 1a and Supplementary Table 1-2) from the ROSMAP cohort^29^, AD PFC samples showed a higher sSNV burden compared to controls with a diagnosis of no or only mild cognitive impairment (Fig. 2c; *p* < 0.01, two-tailed proportion test; OR = 2.1). In a second, independent RNA-seq dataset from the MayoRNAseq project^30^, consisting of 300 brain samples from the temporal cortex and cerebellum of 92 patients who died with neuropathologically confirmed AD and 82 matched controls (Extended Data Fig. 1a and Supplementary Table 1-2), AD temporal cortex samples showed a consistent increase of sSNV burden compared to neurotypical controls (Fig. 2d; *p* = 0.01, two-tailed proportion test; OR = 2.2), with a remarkably similar odds ratio to that seen in the ROSMAP PFC samples. Interestingly, the disease-specific enrichment of sSNV was limited to the temporal cortex samples and not observed in cerebellum (Fig. 2d; *p* = 1, two-tailed proportion test), a brain region not severely affected in AD^36^. The observed greater sSNV burden in AD remained significant after controlling for potential confounding factors including sex, age, RNA-seq coverage, neuronal proportion, and batch effects (Fig. 2e and Extended Data Fig. 1b; *p* = 0.01, linear regression). This enrichment persisted even when only the subset of sSNVs predicted to have deleterious impact on protein function were considered (Extended Data Fig. 1c-d; *p* = 0.047, linear regression).

To ensure that the larger number of somatic mutations in AD brains did not reflect contamination by blood, we measured the presence of blood cell types by analyzing gene markers for blood cells in both bulk and snRNAseq data of ROSMAP and MayoRNAseq (see details in Methods). We confirmed that blood contamination as measured by blood-related transcripts in these brain samples is minimal (Extended Data Fig. 1e); correcting our data for any minimal blood did not change the elevated burden of somatic mutation in AD brains (Extended Data Fig. 1f). Our results from these two RNA-seq datasets consistently suggested that clonal somatic mutations in the cerebral cortex are increased in AD patients.

Using Gene Ontology (GO) annotation, we observed that sSNVs found in AD brains were significantly enriched in genes related to ubiquitin-dependent proteolysis, which has been reported to be associated with AD pathogenesis^37^, as well as in genes that regulate cell cycle and proliferation (adjusted *p* < 0.05, hypergeometric test), and this enrichment pattern was not found in sSNVs identified in control brains (Fig. 2f). Considering the role of proliferation-related genes in amplifying somatic mutations, our results suggested that somatic mutations in proliferation-related genes may be more common in AD cerebral cortex.

### Somatic mutation in proliferation-related genes

As an orthogonal and more sensitive approach to examining the mutational burden in proliferation-related genes in AD, we designed a hybrid capture gene panel covering 149 cancer driver genes with UMI barcoding (Supplementary Table 3), and sequenced DNA from the PFC of 190 AD patients and 121 matched controls from the ROSMAP cohort at an average sequencing depth of >1000X after UMI collapsing (Supplementary Table 4 and Extended Data Fig. 2a-b). By exponentially reducing base-calling errors when generating the consensus sequence from multiple reads derived from the same original DNA molecule, this UMI-based panel sequencing detects somatic mutations with MAFs as low as 0.1% (Extended Data Fig. 2c-d), with much higher sensitivity and precision than previous methods not employing consensus error correction^38^. Using our customized computational pipeline, we successfully identified 199 sSNVs and 13 sIndels that were exclusively present in a single DNA sample (the “stringent” list; Supplementary Table 5). To increase the detection power, we further allowed recurrent mutations when they were specifically enriched in AD or control samples, which expanded our list to 1001 sSNVs and 20 sIndels, respectively (the “sensitive” list; Supplementary Table 5 and Extended Data Fig. 3a-b). The mutation spectrum of sSNVs is consistent with the cell division/mitotic clock signature SBS1 (Extended Data Fig. 3a; cosine similarity 0.92), suggesting that mutations predominantly occurred during cell division. We randomly selected 22 sSNVs with a range of MAFs for validation using amplicon sequencing, along with 17 potentially pathogenic sSNVs identified in AD brains that were predicted to be deleterious, and all of the 10 frameshift sIndels in the “sensitive” list. Thirty-five of 39 (90%) tested sSNVs and 8 of 10 (80%) sIndels successfully validated in newly extracted DNA samples from the corresponding PFC samples, confirming the high accuracy of our somatic mutation calling strategy even for those with MAFs as low as 0.1% (Extended Data Fig. 2e-g).

With similar sequencing depth and coverage between AD and control PFC samples (Extended Data Fig. 2a-b), the stringent pipeline revealed that AD brains harbored significantly more sSNVs among the 149 targeted genes than aged-matched controls (Fig. 3a; *p* = 0.008, two-tailed proportion test; OR = 1.6). When using the sensitive pipeline, which allows recurrent mutations, the sSNV increase in AD brains became even more significant (Fig. 3b; *p* = 0.001, two-tailed proportion test; OR = 1.3), and this pattern remained significant after controlling for confounding factors including sex, age, sequencing coverage, and post-mortem interval (Fig. 3c; *p* = 0.03, linear regression).

**Fig. 3.**
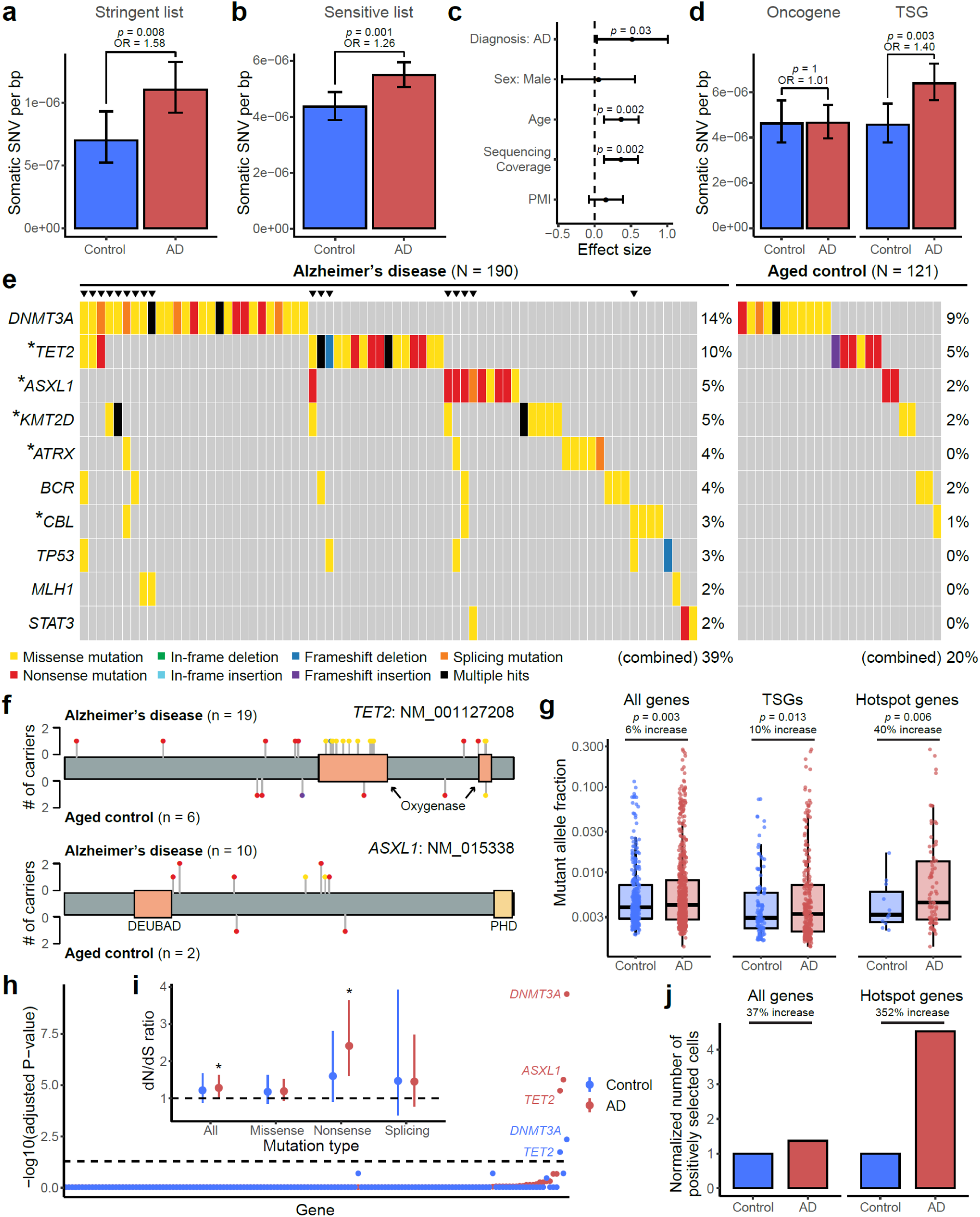
Elevated burdens of somatic mutations in cancer driver genes in AD brains. **a-b**, AD prefrontal cortex samples harbor significantly more sSNVs in 149 targeted cancer driver genes than matched controls, using both the sSNV list of stringent (**a**) and sensitive (**b**) identification pipelines. The sensitive list additionally contains recurrent sSNVs if they were specifically enriched in the AD or control groups. **c**, Linear regression modeling confirmed that the AD effect on greater sSNV burden remains significant (*p* = 0.03) after controlling for potential confounding factors. In addition to AD status, age is also positively correlated with the sSNV burden (*p* = 0.002). **d**, The significant increase of sSNV burden in AD brains was only observed for tumor suppressor genes (TSGs) but not for (proto-)oncogenes. **e**, Top 10 recurrently mutated genes in AD and control brains. Different types of protein-altering sSNV and sIndel are shown in various colors, where “multiple hits” (black) denotes multiple protein-altering mutations in the same gene. Asterisks denote the five “hotspot” genes that contain significantly more somatic mutations in AD patients than matched controls (*p* < 0.05, one-tailed proportion test). Triangles highlight individuals that carry mutations in multiple genes. **f**, Distribution of somatic mutations in two AD hotspot genes, *TET2* and *ASXL1*. The color and height of each lollipop denote the mutation type and the number of carrying individuals. **g**, Somatic mutations in AD brains showed significantly higher allele fractions than controls (two-tailed t-test), with a larger increase when only considering TSGs or AD hotspot genes, suggesting the clonal expansion of cells that carry the somatic mutations. The increase in allele fraction was calculated using the ratio of medians between AD and control groups. Boxplots show median and the first and third quartiles, with whiskers denoting 1.5 * IQR from hinges. **h**, Positive selection of individual genes in AD and control somatic mutations. Y-axis denotes p-value for testing if the gene’s dN/dS ratio is higher than 1, with Benjamini-Hodgberg’s multiple hypothesis testing correction. *DNMT3A*, *ASXL1*, and *TET2* show significant positive selection in AD brains, stronger than in control brains. **i**, dN/dS ratios across all the 149 targeted genes, in which the rates of all protein-altering mutations, missense mutations, nonsense mutations, and splicing mutations are compared with the background neutral rate estimated by synonymous mutations. Asterisks denote p-value < 0.05. **j**, AD brains harbor more positively selected cells than control brains, especially when we only consider somatic mutations in AD hotspot genes. The number of positively selected cells was inferred based on the gene-specific dN/dS ratio, the count of somatic mutation per sample, and the average MAF (see details in Methods). **a**-**d** and **i**, Error bar, 95% CI.

In addition to the increased sSNV in AD, we also found age as an independent factor positively correlated with the sSNV burden (Fig. 3c; *p* = 0.002, linear regression) and the proportion of sSNV carriers (Extended Data Fig. 3c), suggesting a likely age-associated accumulation of somatic mutations in proliferation-related genes in both normal and diseased brains. Previous studies highlighted the age-related accumulation of low-MAF (<1-5%) somatic mutations in cancer driver genes in blood^39^. Our finding about age-related accumulation in brain is consistent with a recent study using deep whole-genome sequencing of a smaller sample^40^, though our study was not designed to specifically test this. We observed that APOE4 carriers tend to have more sSNV than non-carriers in both AD and control groups, though this pattern did not reach statistical significance (Extended Data Fig. 3d; *p* = 0.09, linear regression).

Interestingly, when we divided cancer driver genes into (proto-)oncogenes and tumor suppressor genes (TSGs), we observed a greater sSNV burden in AD for TSGs but not for oncogenes (Fig. 3d). Considering that TSGs lead to proliferation when they are inactivated by loss-of-function mutations throughout the gene body, but oncogenes are usually only activated by specific, recurrent, gain-of-function alleles affecting critical domains, our results suggested that most sSNVs are associated with AD by a loss-of-function of TSGs. Besides sSNV, we also observed more frameshift sIndels in AD brains (5 in AD versus 2 in control; Supplementary Table 5), though this enrichment did not reach significance in this small sample size.

Examination of the mutation burden at the individual-gene level revealed that somatic mutations in the top 10 most commonly mutated genes were found in 39% of the AD patients compared to only 20% of the aged controls (Fig. 3e); brain samples carrying mutations in multiple genes were exclusively found in the AD cohort but not in controls (*p* = 0.0002, hypergeometric test). Five “hotspot” genes—*TET2*, *ASXL1*, *KMT2D*, *ATRX*, and *CBL—*harbored nominally more somatic mutations in AD brains than controls (Fig. 3e; *p* < 0.05, one-tailed proportion test), though these individual gene burdens were not significant after multiple hypothesis testing correction for 149 genes. All “hotspot” genes represent critical TSGs and have been widely implicated in various cancers^41^ and CH^42^. Most AD somatic mutations in *ASXL1* were nonsense mutations broadly distributed across the encoded protein, including two recurrent alleles observed in multiple AD patients, similar to what is seen in *ASXL1* mutations in CH events of blood; AD patients showed missense mutations in *TET2* that clustered in its critical oxygenase domains (Fig. 3f), a similar mutational pattern to that seen in CH (Extended Data Fig. 3e) but not seen in aged controls. Somatic mutations in AD brains showed significantly higher MAFs than did mutations in control brains, especially in the five hotspot genes, where the average MAF was 40% increased, suggesting that many somatic mutations found in AD drive the clonal expansion of cells that carry them to a greater extent than in control brains (Fig. 3g). To further validate this, we examined the signal of positive selection for these mutations and found that somatic mutations in AD brains experienced stronger positive selection in AD brains, evidenced by elevated dN/dS ratios (Fig. 3h-i) as well as a greater abundance of positively selected cell (Fig. 3j). In addition to individual genes, we observed that AD patients had significantly more somatic mutations in PI3K-PKB/Akt pathway genes than controls (Extended Data Fig. 3f; *p* < 0.05, one-tailed proportion test), a pathway that has been previously suggested to be enriched with somatic mutations in AD brains^28^. Overall, our panel sequencing results revealed more frequent somatic mutations in cancer driver genes of AD brains, highlighting their potential roles in driving the clonal expansion of certain proliferating cell types during AD pathogenesis.

### Microglia enrichment of proliferation-related somatic mutation

The overlap of many specific driver genes mutated in AD with those implicated in clonal blood disorders suggested that microglia, which share a very early lineage with peripheral myeloid cells, might be the carrier cells of these mutations in AD brains. To test this, we developed a fluorescence-activated nuclei sorting (FANS) method to specifically isolate microglial nuclei from frozen postmortem brain tissues using an antibody targeting CSF1R (Extended Data Fig. 4a), a well-known cell surface marker for microglia whose nuclear localization and function have been recently reported^43^. Our subsequent snRNAseq (Fig. 4a) and ddPCR (not shown) results confirmed that >75% of sorted nuclei belonged to the microglial cluster in both AD and control brains, verified by expression of microglia marker genes including *CX3CR1*, *TMEM119*, and *P2RY12* (Extended Data Fig. 4b). Interestingly, another 4-9% of the nuclei were classified as CNS-associated macrophages (CAMs; Fig. 4a and Extended Data Fig. 4b), a recently identified class of brain-resident myeloid cells with high expression of *MS4A7* and *MRC1*^44^, while the remaining cells represented scattered neural cells or pericytes. Both microglia and CAMs are brain-resident macrophages predominantly derived from erythromyeloid progenitors during embryogenesis^45^, but recent studies also report a contribution of hematopoiesis-derived immune cells to the brain macrophage pool in adulthood^46,47^.

**Fig. 4.**
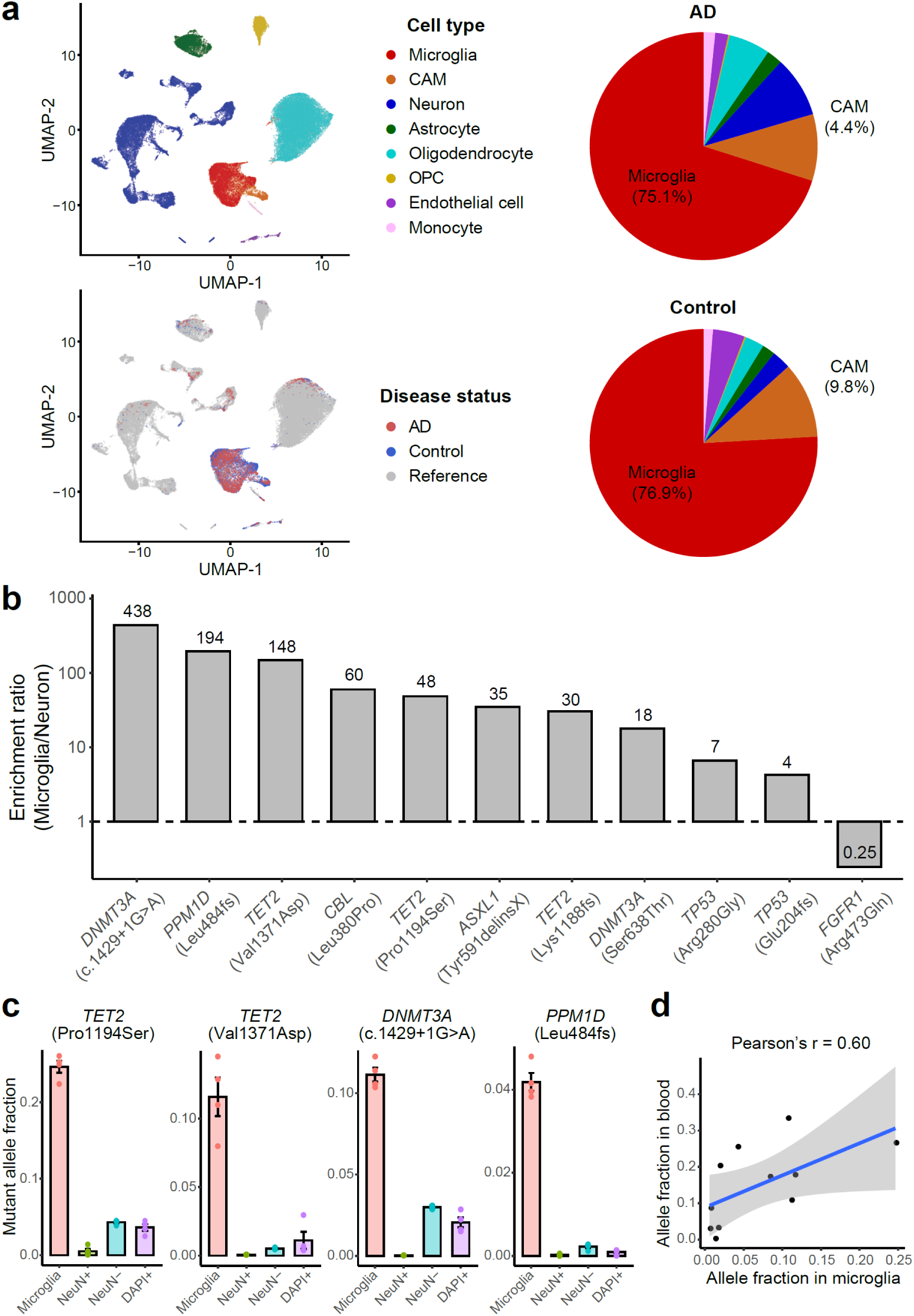
Deleterious somatic mutations are enriched in microglial clones of AD brains. **a**, 10X snRNAseq confirms the high purity and unbiased representation of microglia in CSF1R+ nuclei sorted from AD and control PFC samples. Clustering results suggest about 80% of the sorted nuclei are microglia (red), whereas another 3-9% are CNS-associated macrophages (CAMs, orange). Minimal blood cell contamination is confirmed with up to 1% monocytes and the absence of B cells, T cells, and red blood cells. OPC, oligodendrocyte progenitor cell. **b**, The ratios of mutant allele fractions between sorted microglial and neuronal nuclei of the same AD brains, estimated by amplicon sequencing. Ten of the 11 profiled AD somatic mutations demonstrated at least 4X microglial enrichment. **c**, Four somatic mutations in CH-associated genes as examples show significantly higher allele fractions in microglia than the fractions in the other three populations (*p* < 0.05, two-tailed Wilcoxon test), suggesting their microglial origins. Each nuclei population was sorted four times from each AD brain sample to serve as replicates. Error bar, SE. **d**, All but the *FGFR1* mutations are shared between microglia and whole-blood samples of the same individual, indicating a common origin of these somatic mutations.

We selected 7 sSNVs and 4 sIndels identified from AD brains, all of which were predicted to be deleterious for critical oncogenes or TSGs, and found a marked enrichment of these mutations in the sorted microglial fraction. We measured the MAF of each somatic mutation in four different populations of sorted cells using amplicon sequencing: microglia (CSF1R+), neurons (NeuN+), glia and other nonneuronal cells (NeuN-), and all cells (DAPI+). All ten sSNVs in TSGs were enriched (4- to 438-fold) in microglia when compared to neurons sorted from the same brain sample (Fig. 4b and Extended Data Fig. 4c). For a splicing sSNV in *DNMT3A* (c.1429+1G>A) and two deleterious missense sSNVs in *TET2* (p.Pro1194Ser and p.Val1371Asp), we observed >10% MAFs in microglia, dramatically higher than the MAFs observed in neurons and other mixed cell populations (Fig. 4c; *p* < 0.05, two-tailed Wilcoxon test), suggesting that mutant cells constitute >20% of all microglia in the sample. The last tested sSNV, in the oncogene *FGFR1* (p.Arg506Gln), is a non-recurrent mutation predicted to cause decreased activation of this oncogene, and was not enriched in microglia. Interestingly, this same AD PFC sample harbored a variant in a TSG gene (*DNMT3A* (c.1429+1G>A)) that was almost exclusively present in microglia, suggesting that these two variants originated in different lineages (Extended Data Fig. 4c), but also showing that all tested variants predicted to confer a proliferative advantage were enriched in microglia. Tested mutations were detected in up to 40% of PFC microglia in carrier brains, implying that they provide strong survival and/or proliferative advantages over microglia that do not carry the mutation.

Analysis of matched blood DNA showed that 10 of the 10 mutations enriched in microglia were also present in blood, with a trend towards a positive correlation between MAFs in microglia and blood (*p* = 0.052, Pearson correlation; Fig. 4d and Extended Data Fig. 4d). We confirmed minimal blood contamination in unsorted bulk brains (as measured by RNA-seq analysis) and in the sorted microglial nuclei (Fig. 4a and Extended Data Fig. 4b) as a cause of this shared presence, but our results do not distinguish between a shared lineage, or migration of myeloid or microglial cells into or out of the brain.

### Mosaic chromosome alterations in AD snRNAseq data

To explore the effects of somatic mutations in microglia in Alzheimer’s disease, we utilized a recent high-quality snRNAseq dataset of middle temporal gyrus neocortex samples obtained from AD donors and age-matched controls, the Seattle Alzheimer’s Disease Brain Cell Atlas (SEA-AD). Due to the high degree of transcriptional noise and sparsity within snRNAseq data, there is no tool available to our knowledge that can reliably call sSNVs without matched DNA-seq^48^. However, several methods have been successful at identifying mosaic chromosomal alterations (mCAs), from snRNAseq data^49–51^. Since recurrent mCA has also been associated with CH and other myeloid overgrowth syndromes^10,11^, generally disrupting specific genes also mutated by sSNV, we hypothesized that AD brains would also carry mCA in microglia-CAMs.

We extracted cells that were annotated as microglia-perivascular macrophages (a subtype of CAMs, hereby called microglia-CAMs) or were identified as microglia-CAMs through automatic cell-typing with scType (Extended Data Fig. 5a-b and Supplementary Table 6), and then called microglia-CAM-specific mCAs within SEA-AD using CONICSmat^49^ for all individuals with a consensus clinical diagnosis of AD (n = 31) or healthy, age-matched controls (n = 31) (Supplementary Table 7). We also called mCAs in excitatory neurons (ExNs), astrocytes, oligodendrocytes, or oligodendrocyte precursor cells (OPCs) and retained only mCAs that were not called in any of these other cell types from the same donor and which passed several stringent filtering criteria (Materials and Methods and Extended Data Fig. 5c).

AD brains harbored nominally more mCAs (4 in AD versus 1 in control; Fig. 5a) and nominally 8-fold more mCA-carrying microglia-CAMs (Fig. 5b; *p* = 0.06, permutation test), though as expected, the SEA-AD sample size was too small for these differences to reach statistical significance. When we analyzed microglia and CAM separately, we observed a stronger trend in microglia than CAMs (Fig. 5c; *p* = 0.07 and 0.11, permutation test). We also observed an increasing trend of mCA in AD individuals versus controls in astrocytes, but not in oligodendrocytes, OPCs, and ExNs (Fig. 5c and Supplementary Table 7), perhaps relating to the widespread astrogliosis reported in AD^52^.

**Fig. 5.**
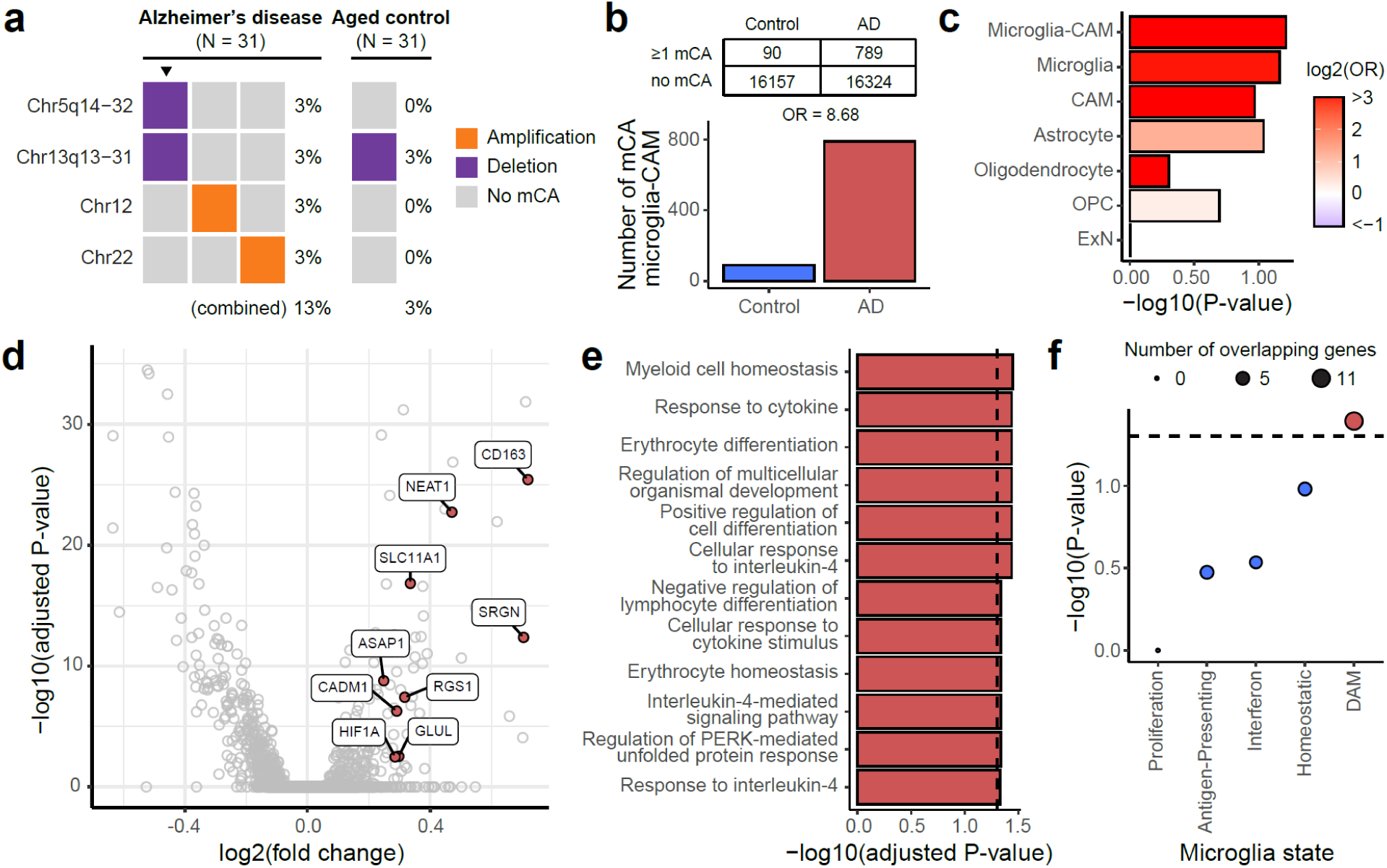
mCAs in AD microglia are associated with a pro-inflammatory, disease-related signature. **a**, Microglia from AD brains contain nominally more mCAs associated with hematopoietic overgrowth syndromes compared to age-matched controls, even in this small sample (N = 31 each). Triangles highlight an individual with multiple mCAs. **b**, AD brains show a trend (*p* = 0.06, permutation test) towards a higher fraction of mCA-carrying microglia than age-matched controls. **c**, Odds ratios of mCA-carrying cells between AD and control individuals across different cell types. Microglia-CAM (*p* = 0.06) and microglia (*p* = 0.07) have the smallest nominal p-values in permutation test compared to CAMs (*p* = 0.11), astrocytes (*p* = 0.09), oligodendrocytes (*p* = 0.50), OPC (*p* = 0.40), and ExN (*p* = 0.99). OPC, oligodendrocyte progenitor cell. ExN, excitatory neuron. **d**, Volcano plot shows differentially expressed genes between AD donor microglia-CAMs with and without mCA. Positive fold-change indicates upregulation in microglia-CAMs with mCA. DAM-associated upregulated genes are colored red. **e**, Significantly (adjusted *p* < 0.05, hypergeometric test) enriched gene ontology terms for genes upregulated in microglia-CAMs with mCA. **f**, Enrichment of microglial state modules^53^ among genes upregulated in microglia-CAMs with mCA. Significant enrichments implicate inflammation and the DAM transcriptional state.

### Transcriptional effect of somatic mutations in AD microglia

While the SEA-AD sample size is too small to demonstrate independent enrichment of mCA in microglia, they are certainly consistent with this, and allowed analysis of the transcriptional effects of mCA in microglia, by creating an integrated snRNAseq atlas of microglia-CAMs identified across AD cases and controls (Extended Data Fig. 6) and identifying differentially expressed genes (DEGs) between mutant and wild-type microglia-CAMs from mCA-carrying AD brains (Fig. 5d and Supplementary Table 8). Using gene ontology (GO) enrichment analysis, we found that DEGs with increased expression in mutant microglia were enriched (adjusted *p* < 0.05, hypergeometric test) for several terms related to immune activation and signaling, suggesting that mutant microglia may upregulate pro-inflammatory pathways (Fig. 5e and Supplementary Table 8).

A recent study identified transcriptional signatures of microglial states in human stem-cell differentiated microglia that emerge in response to various CNS challenges, such as apoptotic neurons, amyloid-beta fibrils, and myelin debris^53^. We used these signatures to further characterize the microglial state associated with mCAs. Using a hypergeometric test for enrichment, we found marginally significant overlap between DEGs that are upregulated in mutant microglia and genes associated with the DAM state (Fig. 5f and Supplementary Table 8; *p* = 0.04). DAMs are specifically enriched in AD brains and have been posited to play a role in modulating the neuroinflammatory response to neurodegeneration^3,54^, suggesting that microglia with mCA may share a similar phenotype in AD.

## Discussion

Our results from three independent AD cohorts, using three orthogonal approaches, revealed a consistently greater burden of somatic mutations in AD cerebral cortex samples when compared to matched controls, suggesting that brain somatic mutation is associated with AD. These somatic mutations were enriched in proliferation-related genes that have been widely implicated in cancer and pre-cancerous conditions, with higher MAFs and stronger positive selection in AD brains, implying their roles in clonal expansion of mutant cells. This was also supported by the enrichment of AD cases with multiple CH-associated sSNVs. We further confirmed that many mutations were specifically present in microglia, and potentially CAMs. Finally, using snRNAseq analysis we found that microglia carrying mCAs associated with clonal overgrowth syndromes showed pro-inflammatory and disease-associated transcriptional signatures compared to wild-type counterparts. While we cannot formally rule out that clonal expansion of mutant microglia represents only a secondary response to proliferative signals in AD brain, the DAM-related signature associated with mCA resembles effects of CH mutations in blood myeloid cells that increase the risk of myocardial infarction and stroke while activating immune cascades including IL1ß, IL6, and others^56^. These similarities suggest analogous roles of microglial mutations in AD that would likely promote neuronal degeneration^57^.

Two recent studies correlating CH mutations in blood with AD risk found no effect^58^ or a surprising protective effect of blood CH on AD^59^. Although many methodological differences exist between those blood studies and our brain study (Supplementary Discussion), the varying results highlight the complexity and limitations of our current understanding of the relationship between myeloid cells and microglia. Bouzid *et al.*^59^ and we both found that microglial driver mutations were typically shared in the blood of the same individual, as did a small earlier study that also found cancer driver mutations in AD brain^27^. Since somatic driver mutations that lead to blood cancer, when dated by lineage analysis, often arise before birth^60^, MiCE mutations may occur in early progenitors of microglial and blood lineages. Under this assumption, microglia carrying the same driver mutations may clonally expand in brain independently from blood. Alternatively, recent studies show that myeloid cells from blood can enter the brain when there is dysfunction of the blood-brain barrier (BBB), an early feature of AD^61^, and can differentiate into microglia-like cells^62^. Other*s* have reported that monocytes can enter the brain and form microglia-like cells even independent of BBB disruption^46,47^. Thus, BBB changes may be a critical feature that might promote access of mutant myeloid cells to the CNS. Conversely, activated microglia can form perivascular clusters in neurodegeneration as a result of BBB breakdown^63,64^ which might allow mutant brain microglial cells access to enter the bloodstream.

Our results suggest that microglia are the major cell type carrying somatic driver mutations. Although our FANS results cannot completely exclude CAMs also carrying these somatic mutations, our CSF1R+ cell population contained 3% and 9% CAMs in AD and control brains, respectively (Fig. 4a), and 5 of the 11 somatic mutations represented >10% cell fractions in the sorted microglial nuclei of AD brains, including the *TET2* p.Pro1194Ser variants with >40% cell fraction. This high MAF seems inconsistent with the mutation being limited to blood-derived macrophages even assuming all CAMs came from the blood myeloid lineage.

Our analysis highlighted five hotspot genes as well as the PI3K-PKB/Akt pathway (including a *PIK3CA* p.His1047Leu activating mutation and three loss-of-function mutations in *TSC1/2*) that were recurrently disrupted by somatic mutations in AD brains. Drugs targeting such genes have been widely used to treat cancer^65,66^, thus they might serve as potential therapeutic agents to suppress somatic-mutation-activated microglia and ultimately neurodegeneration in AD. Since the role of disease-associated microglia in neuronal loss and dysfunction may be a common feature shared across many neurodegenerative diseases as well as in age-associated cognitive decline, studying somatic mutation in AD may provide an important new approach to understanding the pathogenic mechanisms of dementia and other neurodegenerative conditions.

## Acknowledgments

We thank B. Stevens, B. Hyman, Po-Ru Loh, and B. Yankner for constructive discussions and suggestions on the manuscript, and R. S. Hill, J. E. Neil, D. Gonzalez, M. Chin, and T. Dolbeare for their help. R. Mathieu and T. Berisha from the Boston Children’s Hospital Flow Cytometry Core and IDDRC Molecular Genetics Core helped with sorting. We thank the donors of postmortem tissues for their invaluable contributions to the advancement of science. The results published here are partly based upon data generated by the TCGA Research Network: https://www.cancer.gov/tcga. This work was supported by R56 AG079857 (A.Y.H., C.A.W., E.A.L.); Alzheimer’s Association Research Fellowship (A.Y.H.); PRMRP Discovery Award W81XWH2010028 (Z.Z.); Edward R. and Anne G. Lefler Center Postdoctoral Fellowship (Z.Z.); T32 GM007753 (M.T.); T32 GM144273 (M.T.); K08 AG065502 (M.B.M.); T32 HL007627 (M.B.M.); Brigham and Women’s Hospital Program for Interdisciplinary Neuroscience through a gift from L. and T. Rand (M.B.M.); Alzheimer’s Disease Research program of the BrightFocus Foundation A20201292F (M.B.M.); Doris Duke Charitable Foundation Clinical Scientist Development Award 2021183 (M.B.M.); The Manton Center Pilot Project Award and Rare Disease Research Fellowship (B.Z.); R25 NS065743 (S.K.); U19 AG060909 (E.L.); Suh Kyungbae Foundation (E.A.L.); DP2 AG072437 (E.A.L.); R01 NS032457-20S1 (C.A.W.); R01 AG070921 and AG078929 (C.A.W., E.A.L.); F-Prime Foundation (C.A.W.); Allen Discovery Center program, a Paul G. Allen Frontiers Group advised program of the Paul G. Allen Family Foundation (C.A.W., E.A.L.). ROSMAP is supported by P30 AG10161, P30 AG72975, R01 AG15819, R01 AG17917, U01 AG46152, U01 AG61356 (D.A.B.). C.A.W. is an Investigator of the Howard Hughes Medical Institute.

## Author contributions

A.Y.H., Z.Z., M.B.M., E.A.L., and C.A.W. conceived and designed the study. A.Y.H. developed RNA-MosaicHunter and performed somatic mutation calling from bulk RNA-seq data, with the assistance of B.Z., D.K., and J.C.. Z.Z. designed and performed panel and amplicon sequencing for bulk brain and sorted cell populations, with the assistance of M.B.M., B.C., L.E., I.R., E.S., S.K., and J.G.. A.Y.H. performed somatic mutations calling on panel sequencing and amplicon sequencing data, with the assistance of J.K.. M.T. performed somatic mutations calling from snRNAseq data and downstream transcriptomic analysis, under the guidance of A.Y.H.. P.L.J. and D.A.B. provided brain tissues and genomic DNA for ROSMAP samples. K.T., M.G., R.H., E.K., and E.L. provided SEA-AD snRNAseq data. A.Y.H, E.A.L., and C.A.W. supervised the study. A.Y.H., Z.Z., M.T., E.A.L., and C.A.W. wrote the manuscript.

---

**Extended Data Fig. 1.**
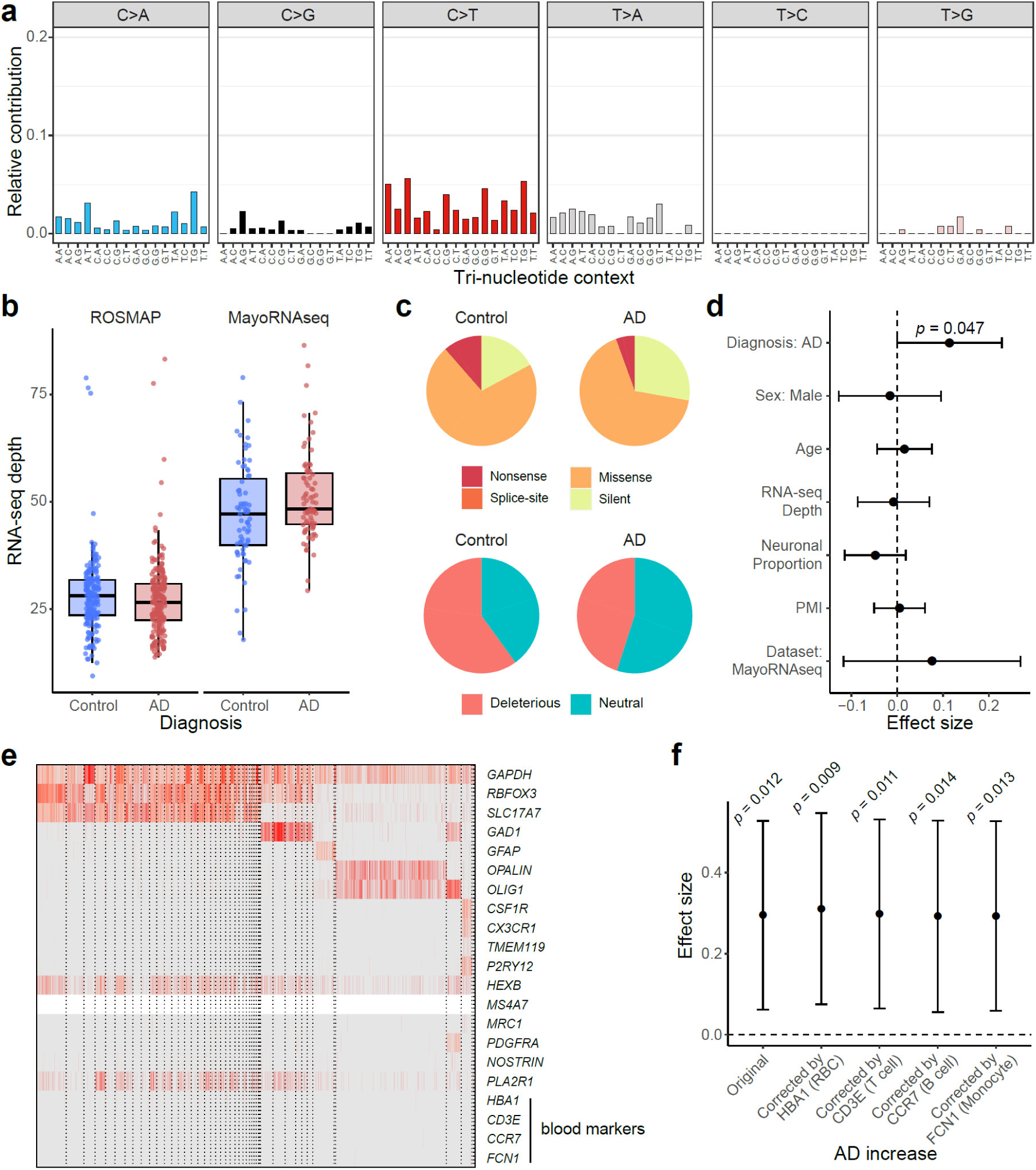
Identification and functional annotation of sSNVs in RNA-seq data. **a**, Mutation type and tri-nucleotide context of sSNVs. T-to-C (A-to-G) candidates were ignored because they were more likely to be RNA-editing sites widespread in the human genome. **b**, Similar sequencing depth between the AD and control brain samples in each AD cohort. The overall higher depth in MayoRNAseq may explain the higher base-line mutation burden in control brain samples than ROSMAP. Boxplots show median and the first and third quartiles, with whiskers denoting 1.5 * IQR from hinges. **c**, Genic annotation and functional impact prediction of sSNVs identified from AD and control brain samples. **d**, AD brains had significantly more deleterious sSNVs than controls (*p* = 0.047, linear regression) after controlling for potential confounding factors. **e**, Absent expression of blood marker genes in snRNAseq of unsorted ROSMAP brains confirmed minimal blood contamination. **f**, The AD increase was consistently significant when the proportion of blood cell types indicated by the expression of marker genes was additionally considered in the linear regression model. RBC, red blood cell. **d**,**f**, Error bar, 95% CI.

**Extended Data Fig. 2.**
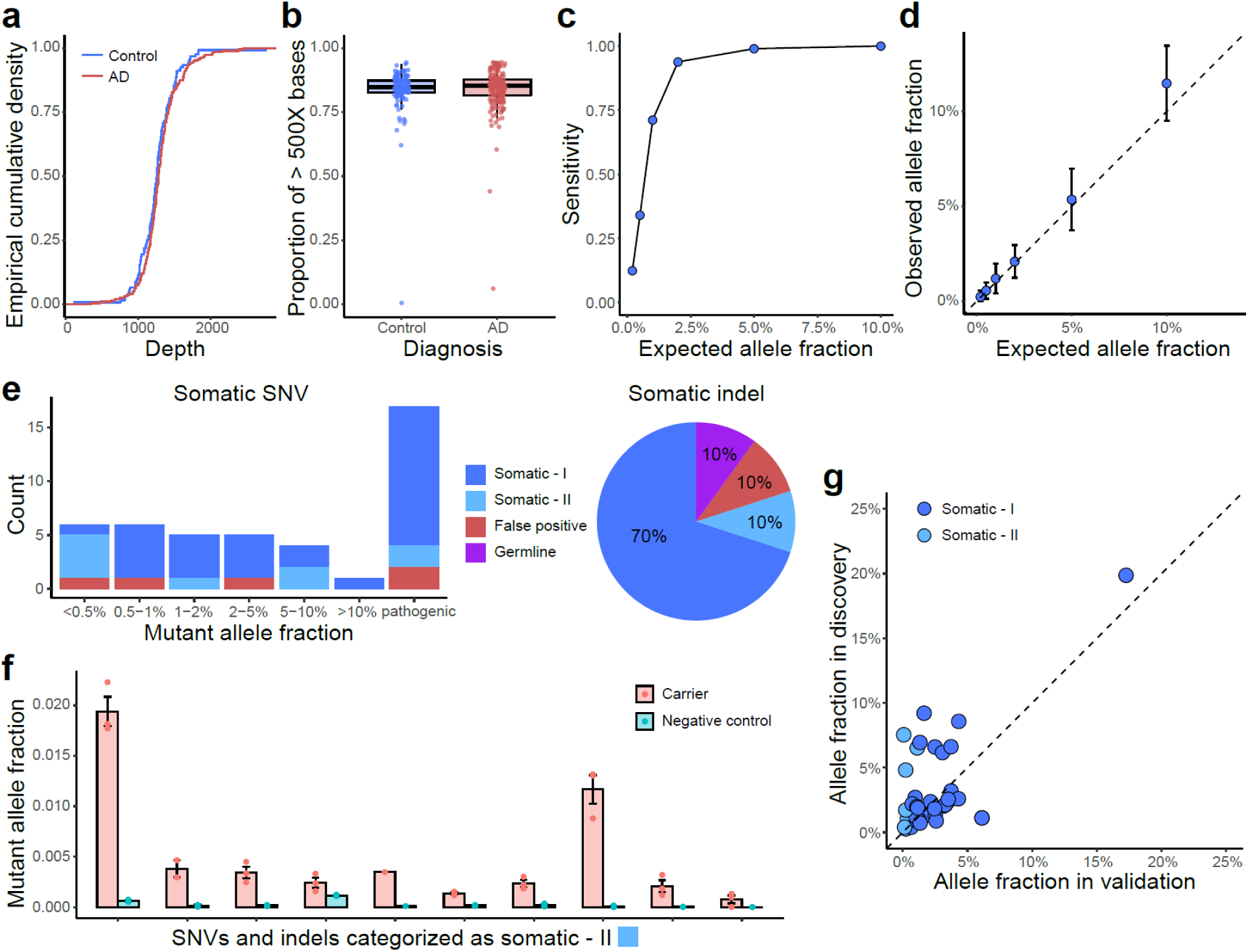
Benchmarking and validation results of sSNVs and sIndels identified from panel sequencing. **a-b**, Comparable sequencing depth (**a**) and coverage (**b**) between AD and control PFC samples, calculated based on the consensus reads after UMI-based read collapsing. **c-d**, Detection sensitivity (**c**) and accuracy of allele fraction estimation (**d**) for our panel sequencing and somatic mutation identification pipeline, benchmarked by *in vitro* mixture of the DNA samples of two unrelated individuals with varied genome ratios. Error bar, SD. **e-f**, Amplicon sequencing validation confirmed high accuracy for identified sSNVs and sIndels in AD and control samples (**e**). Somatic-I mutations are those with mutant allele fractions at least 3X larger than the fractions of the other two error alleles of the same genomic position, whereas somatic-II are those that were further validated by comparing their mutant allele fractions in a negative control sample (**f**). Error bar, SE. **g**, Mutant allele fraction of validated somatic mutations between panel sequencing (discovery) and amplicon sequencing (validation). Amplicon sequencing was performed using newly extracted DNA from the corresponding brain sample, therefore the allele fractions could be varied between the discovery and validation stages.

**Extended Data Fig. 3.**
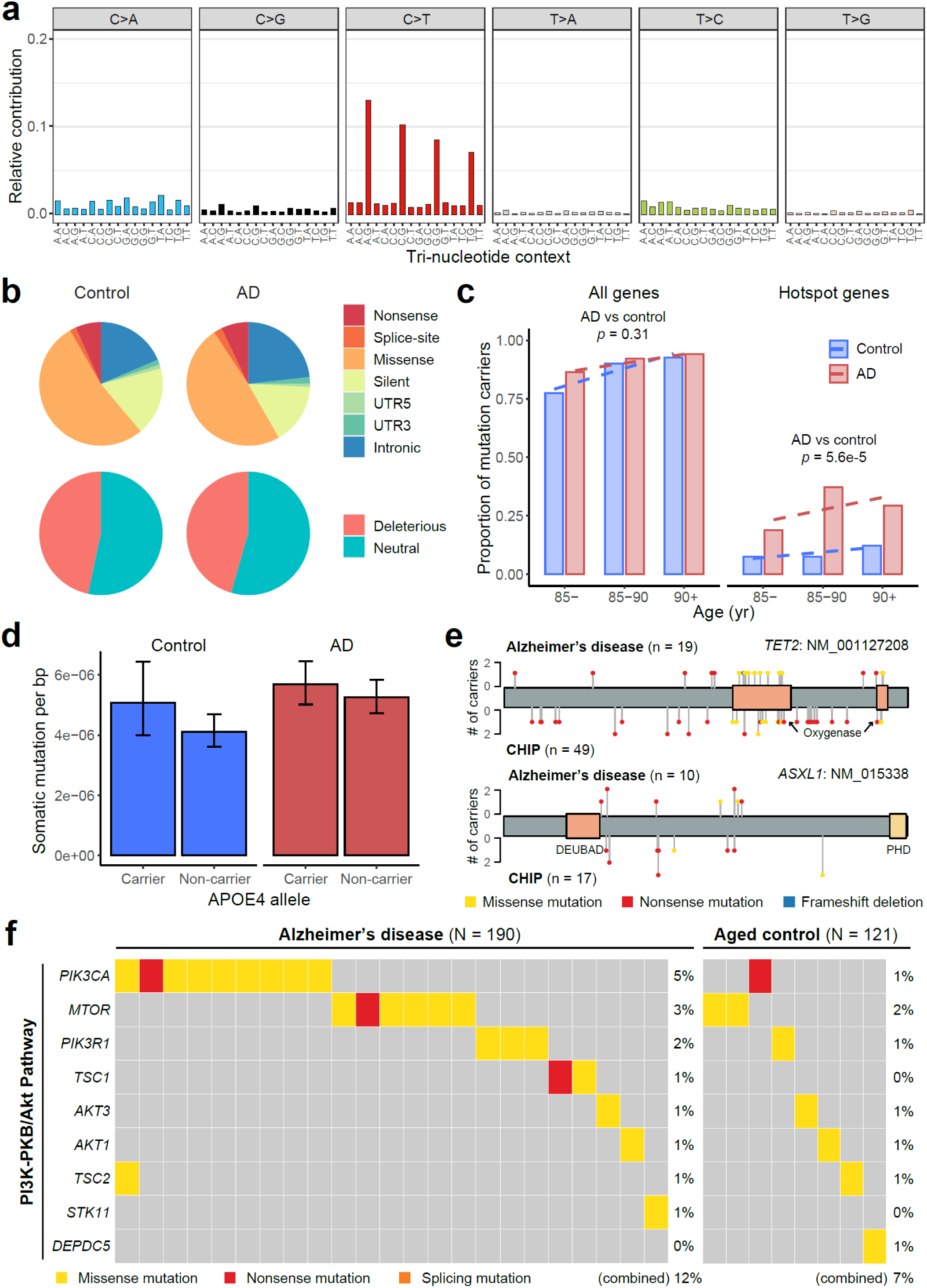
Identification and functional annotation of sSNVs in panel sequencing data. **a**, Mutation type and tri-nucleotide context of sSNVs. **b**, Genic annotation and functional impact prediction of sSNVs identified from AD and control PFC samples. **c**, The proportion of somatic mutation carriers increases with age. AD patients had a significantly larger proportion of carriers with somatic mutations in AD hotspot genes than matched controls (*p* = 5.6e-5, linear regression). **d**, APOE4 carriers tend to have higher burden of sSNVs than non-carriers in both AD and control groups (*p* = 0.09, linear regression). **e**, Similar distributions between somatic mutations identified in AD brains and previously reported CH-associated mutations in blood. **f**, Genes in the PI3K-PKB/Akt pathway contained significantly more somatic mutations in AD brains (12% of AD samples vs 7% of control samples; *p* < 0.05, one-tailed proportion test).

**Extended Data Fig. 4.**
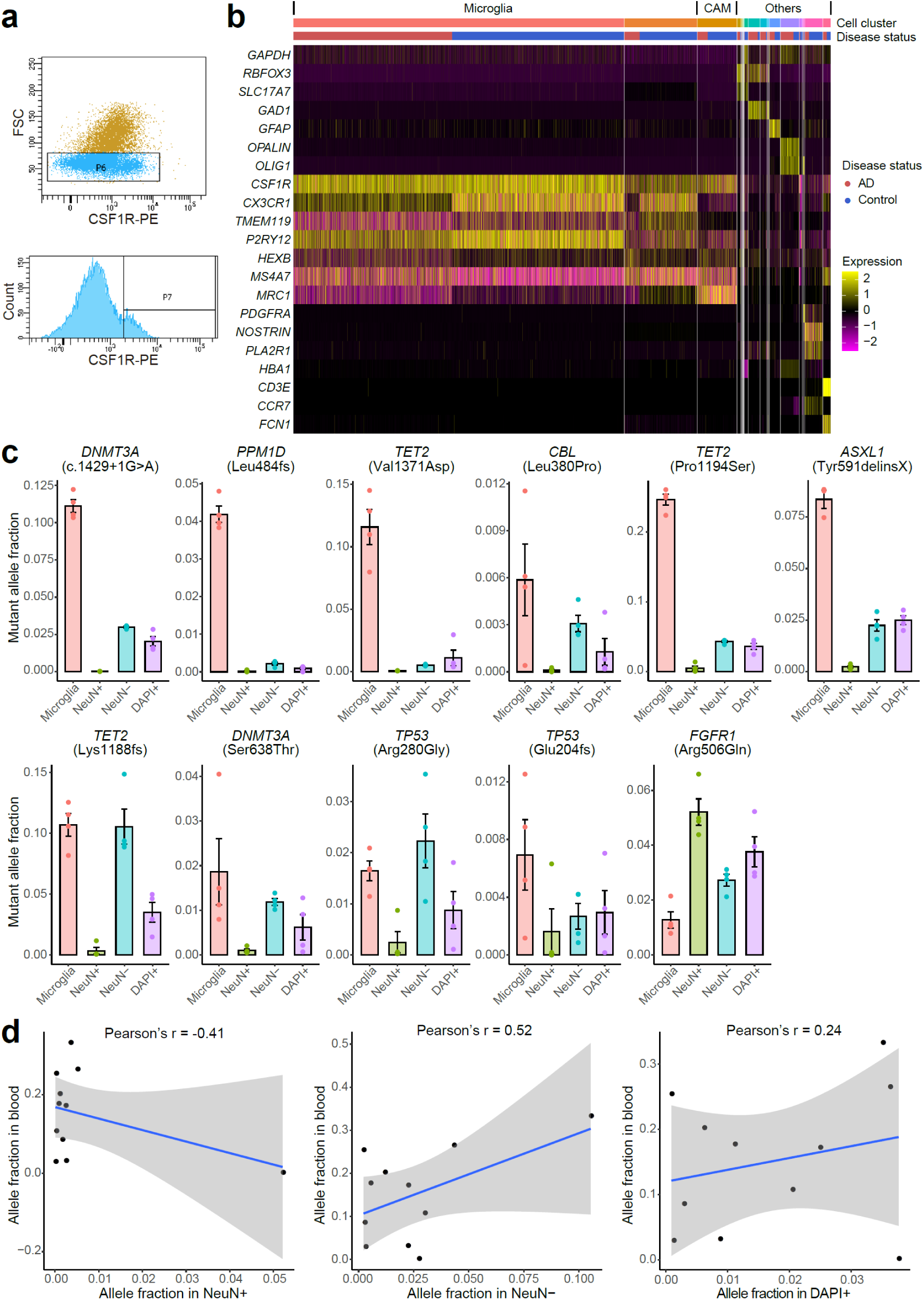
Microglial purity and mutant allele fraction of FANS-sorted nuclei population. **a**, Selectively isolated microglia from frozen brain tissues using FANS with an antibody targeting epitopes of CSF1R, a gene highly expressed in microglia. **b**, Marker gene expression profile for 10X single-nucleus RNA-seq of CSF1R+ sorted nuclei. Each column represents a single nucleus, clustered by PCA based on their expression similarity. About 75-77% of the sorted nuclei are microglia with high expression of *CX3CR1*, *TMEM119*, and *P2RY12*, whereas another 4-9% are CNS-associated macrophages (CAMs). Markers for blood cell types (*HBA1*: red blood cell; *CD3E*: T cell; *CCR7*: B cell; *FCN1*: monocyte) confirm the minimal presence of blood cells in sorted nuclei. CNS, central nervous system. AD microglia showed generally reduced expression of *CX3CR1* and *P2RY12*, consistent with previous findings in AD^3^. **c**, Mutant allele fractions across different sorted nuclei populations for all the 11 profiled AD somatic mutations. Four mutations are shown in Fig. 4c as examples. In all but the *FGFR1* mutation, we observed significantly higher allele fractions in microglia than in neurons (NeuN+). Each population of nuclei was sorted four times from each AD brain sample to serve as replicates. Error bar, SE. **d**, The correlation of mutant allele fractions between blood and three nuclei populations (NeuN+, NeuN-, and DAPI+) sorted from matched brain samples.

**Extended Data Fig. 5.**
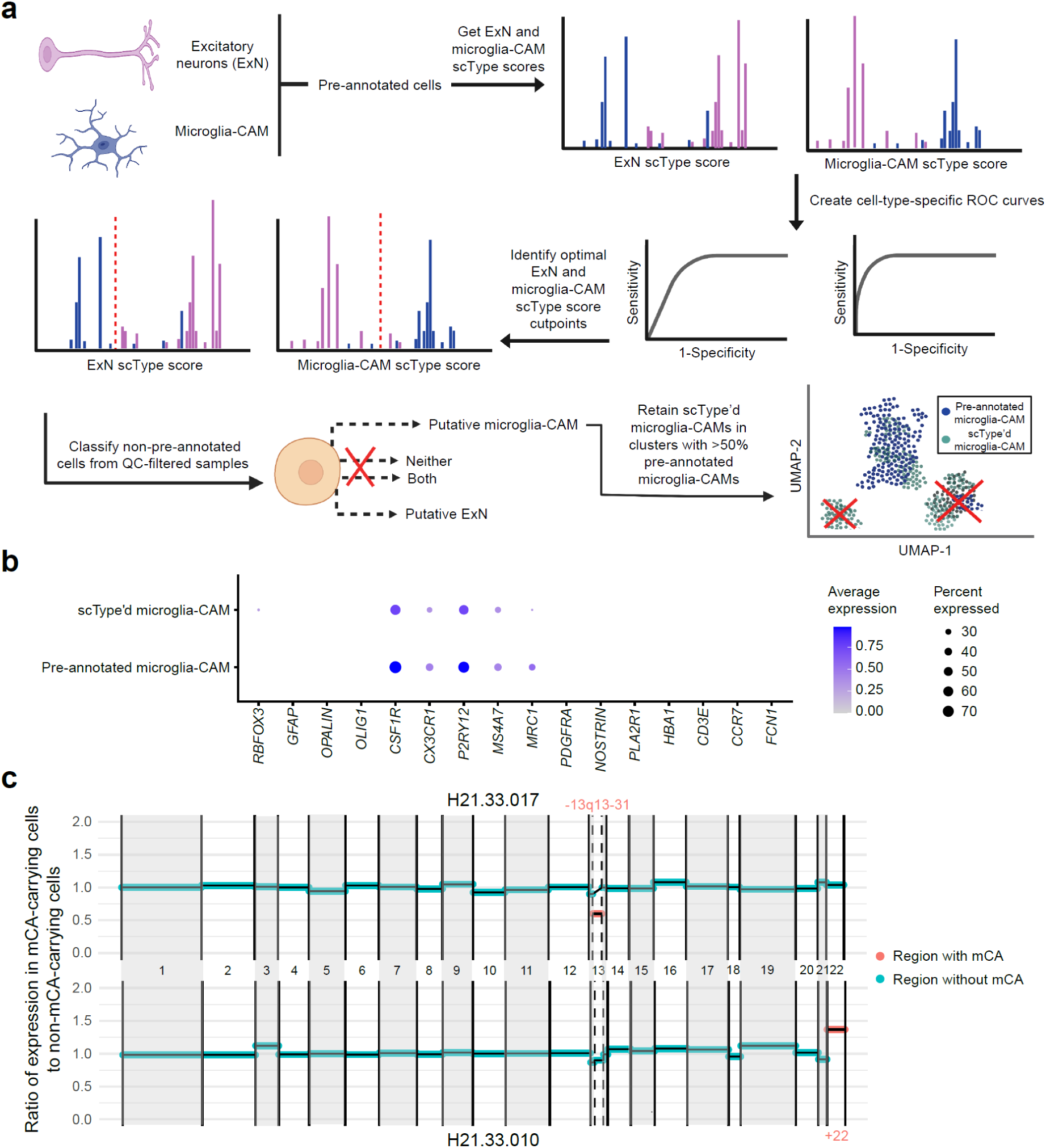
mCA burden analysis in microglia-CAMs and identification of additional microglia-CAMs with scType. **a**, Schematic representation of supervised learning framework and quality-control metrics used to detect additional high-quality microglia-CAMs from SEA-AD. **b**, scType’d and pre-annotated microglia-CAMs show similar marker gene expression profiles, with specific expression of microglia and CAM marker genes. **c,** Examples of mCA called in two AD individuals, H21.33.017 (chr13p13-31 deletion) and H21.33.010 (chr22 amplification). Normalized median ratio of expression in mCA-carrying cells versus non-carrying cells displayed per chromosomal region, with chromosome size proportional to number of expressed genes in microglia-CAMs from that chromosome.

**Extended Data Fig. 6.**
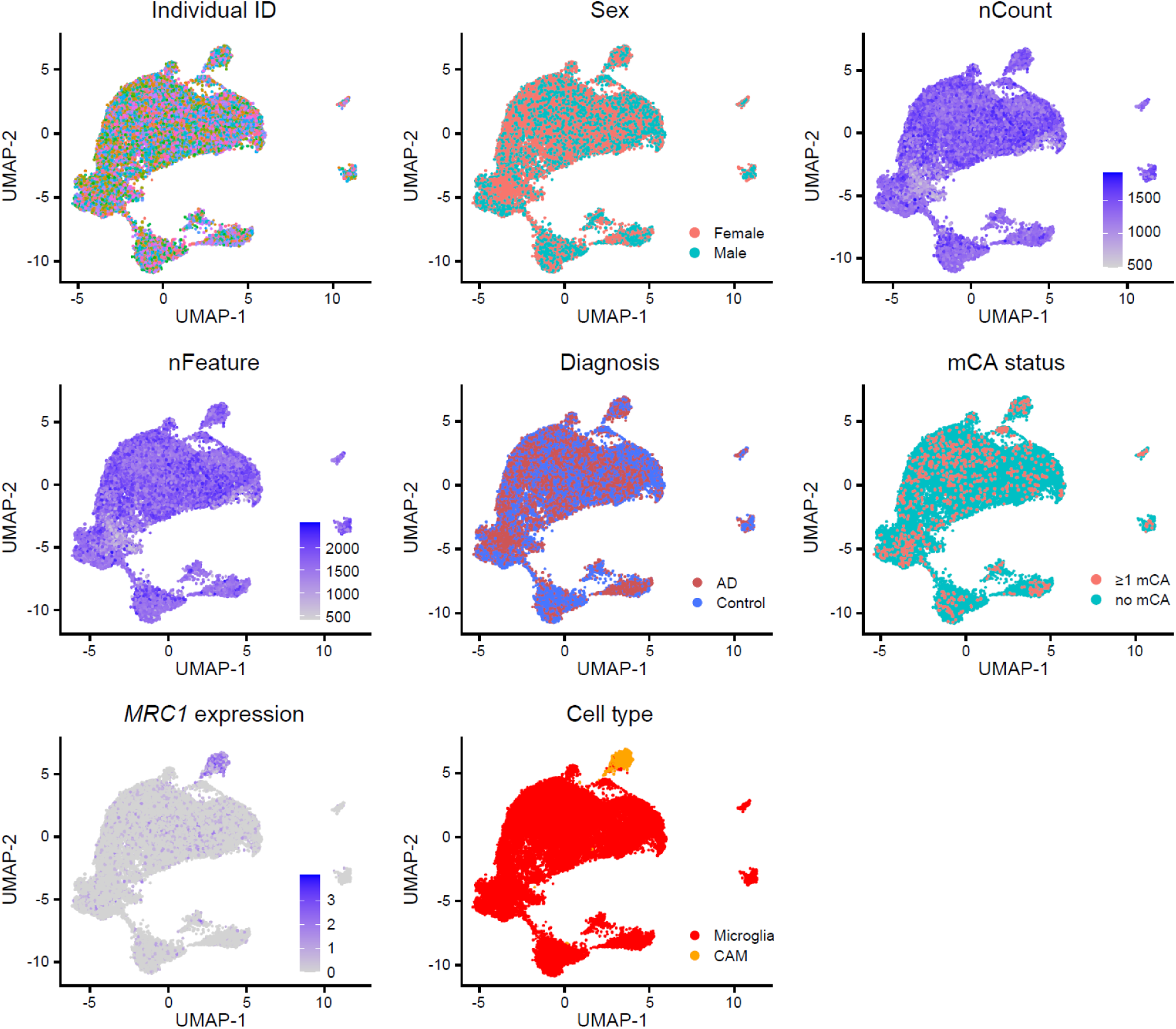
Integrated snRNAseq atlas of microglia-CAMs in AD and healthy controls. UMAP visualization of covariates of interest does not reveal significant clustering by individual ID, nFeature, or nCount, consistent with successful integration across samples. Microglia and CAMs (with high *MRC1* expression) separate into distinct clusters.

**Extended Data Fig. 7.**
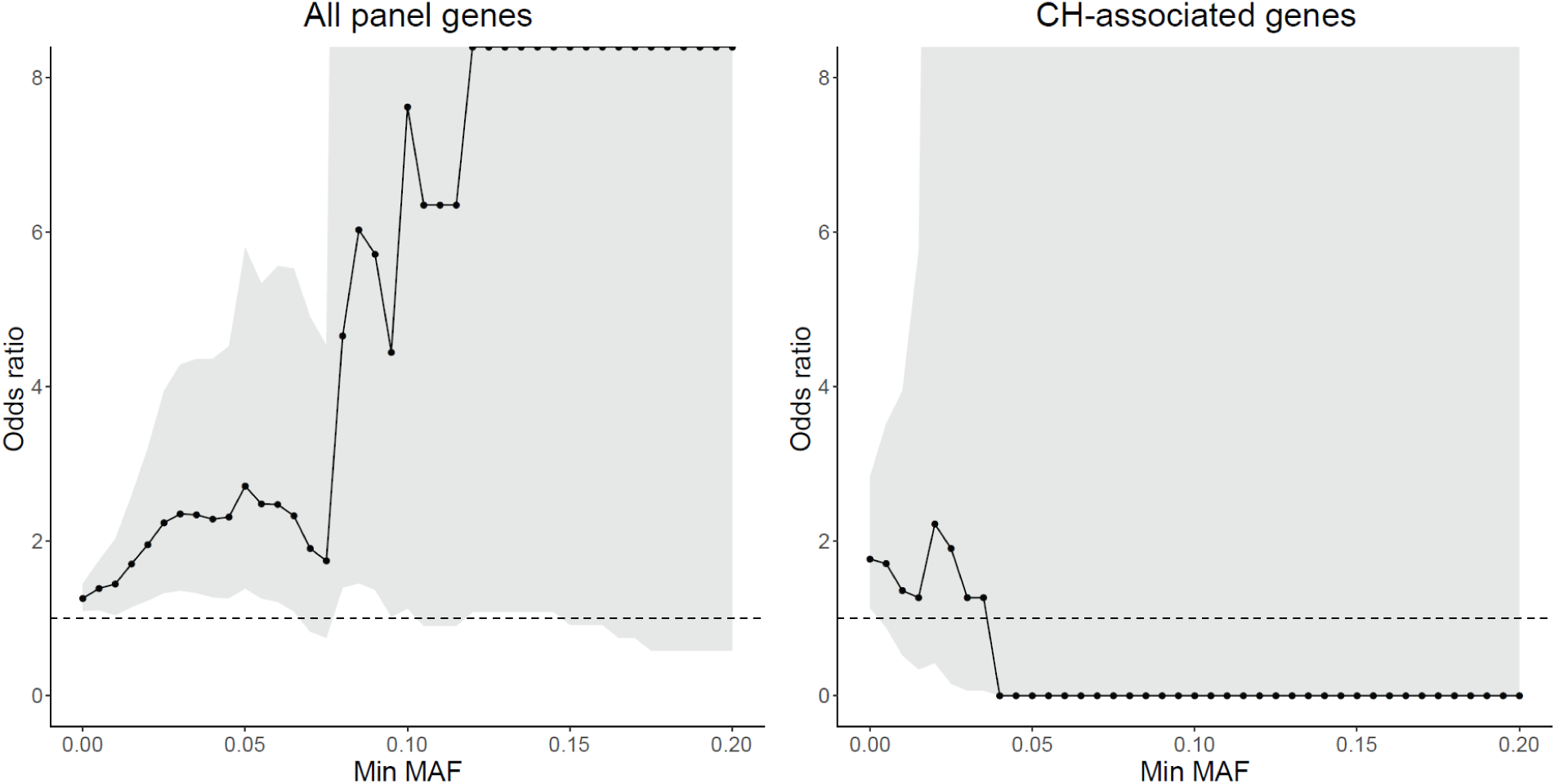
The odds ratio of AD enrichment for sSNVs with different MAF cutoffs. When we consider all the 149 genes targeted by the panel sequencing, we observe a consistent trend of AD enrichment even for sSNVs with 5% or more MAF. In comparison, when we only consider deleterious somatic mutations in CH-associated genes, the odds ratio becomes smaller than 1 when MAF is larger than 4% though with a very large confidence interval. The dashed line represents the odds ratio of 1, and odds ratios larger and smaller than 1 denote the enrichment and depletion of sSNV in AD, respectively.

## Captions for online supplementary tables

**Supplementary Table 1. RNA-seq sample information and summary.** PMI, post-mortem interval.

**Supplementary Table 2. sSNV candidates identified from RNA-seq samples.** sSNVs of ROSMAP and MayoRNAseq samples are listed in separate tabs.

**Supplementary Table 3. List of 149 cancer driver genes in panel sequencing.** TSG, tumor suppressor gene.

**Supplementary Table 4. Panel sequencing sample information and summary.** PMI, post-mortem interval.

**Supplementary Table 5. sSNV and sIndel candidates identified from panel sequencing samples.** sSNVs and sIndels called by the stringent and sensitive pipelines are listed in separate tabs.

**Supplementary Table 6. snRNAseq sample and cell-type annotation information and summary.**

**Supplementary Table 7. mCA candidates identified from snRNAseq samples.**

**Supplementary Table 8. Differential expression and functional annotation results between mutant and wild-type microglia-CAMs from mCA-carrying AD individuals.** Pct.1, expression in microglia-CAM carrying mCA. Pct.2, expression in microglia-CAM that do not carry mCA.

## Methods

### Sample information

Our study involves samples and sequencing data from three large-scale Alzheimer’s disease (AD) studies, ROSMAP, MayoRNAseq, and SEA-AD. The ROSMAP study consists of two prospective studies of aging, The Religious Order Study (ROS) and the Memory and Aging Project (MAP), in which the participants were enrolled by the Rush Alzheimer’s Disease Center with detailed cognitive and neuroimaging phenotyping as well as structured neuropathologic examination during the autopsy at the time of death^1^. The MayoRNAseq study performed detailed clinical phenotyping and multi-omic profiling for 278 participants collected by the Mayo Clinic Brain Bank and Banner Sun Health Research Institute^2^. The SEA-AD study performed single-cell multi-omics, quantitative neuropathology, and deep clinical phenotyping on post-mortem brain tissue from 84 aged donors and 5 additional younger neurotypical controls collected at the University of Washington BioRepository and Integrated Neuropathology laboratory and Precision Neuropathology core. Postmortem samples in all studies were collected and de-identified following the protocol of the corresponding Institutional Review Board with informed consent. The diagnosis of AD was based on the consensus conclusion from all postmortem data generated by neurologists with expertise in dementia and neurodegeneration.

The RNA-seq bam file and the vcf file of germline mutation calls from matched whole-genome sequencing data generated by the ROSMAP and MayoRNAseq studies were downloaded from the AMP-AD Knowledge Portal, along with the detailed demographic and clinical information for each sample. The raw single-nucleus RNA sequencing (snRNAseq) .h5 matrices for SEA-AD and corresponding clinical and technical metadata were also downloaded from AMP-AD Knowledge Portal. Supplementary Table 1 and 6 summarized all the bulk and single-nucleus brain RNA-seq samples analyzed for somatic mutation calling. The ROSMAP dataset consists of the prefrontal cortex (PFC) samples of 228 AD patients and 338 age-matched controls with no or mild cognitive impairment collected by the ROSMAP project. The MayoRNAseq dataset consists of the temporal cortex and cerebellum samples from 92 AD patients and 82 age-matched controls collected by Mayo Clinic, most of whom have RNA-seq from both the temporal cortex and cerebellum samples. The SEA-AD dataset consists of the middle temporal gyrus of temporal cortex from 31 AD patients and 32 age-matched controls. In each dataset, the AD and control samples showed similar distributions in sex, age, post-mortem interval, and sequencing depth (Supplementary Table 1 and 6).

In addition to access to the sequencing data, we obtained genomic DNA (gDNA) from 190 AD patients and 123 controls without cognitive impairment from ROSMAP for panel sequencing (Supplementary Table 4), though this donor list has minimal overlap with the donor list of the brain RNA-seq dataset due to the limited sample availability. Additional dorsolateral PFC brain samples and gDNA from peripheral blood samples were also obtained from ROSMAP to confirm the presence of somatic mutation and further study the cell type identity of mutation-carrying cells.

### Design of RNA-MosaicHunter

Compared to DNA-seq data, RNA-seq data has unique features that need to be addressed for somatic mutation calling. First, the exon-intron structure in mRNA requires the spliced alignment of RNA-seq reads onto the human reference genome, which increases the chance of alignment errors when the overhang sequence is relatively short^3^. Second, the widespread adenosine-to-inosine (A-to-I) RNA editing sites across the human genome^4^ are indistinguishable from A-to-G somatic mutations in RNA-seq data, because inosine will be recognized as guanine (G) in Illumina sequencers. Third, the allele-specific expression^5^, a phenomenon that the paternal and maternal alleles have different expression levels, is observed in many autosomal and X chromosome genes, which can lead to deviated allele fraction estimation in RNA-seq data.

To address these technical issues, we developed RNA-MosaicHunter, which was derived from MosaicHunter^6,7^, a bioinformatic tool designed to identify somatic single-nucleotide variants (sSNVs) in DNA-seq data. RNA-MosaicHunter consists of two major components, a Bayesian genotyper to distinguish real mutations from base-calling errors, followed by a series of empirical error filters to remove artifacts introduced from various sources (Fig. 1a). In the Bayesian genotyper, *G* denotes the genotype state, π denotes the prior probability of each genotype inferred from the population mutant allele fraction p_alt_ and default somatic mutation rate p_m_, and *d, **q**,* and ***o*** denote the depth, base qualities, and bases for calculating genotype likelihoods from the observed sequencing data. Since the mutant allele fraction in RNA-seq data can be affected by allele-specific expression, we considered the posterior probability of both germline heterozygous mutation and somatic mutation in our list of mutation candidates for subsequent error filters, and further distinguished somatic mutations from germline heterozygous mutations by using the genotyping results from matched whole-genome or whole-exome sequencing data obtained from the same individual. In addition, RNA-MosaicHunter also incorporated other filters to exclude 1) candidates with less than 5% mutation allele fraction or less than 5 mutant-supporting reads; 2) candidates that are in repetitive and homopolymer regions; 3) candidates that have a significant bias in strand, mapping quality, or within-read position between the reference and mutant alleles; 4) candidates that show complete linkage to adjacent candidates on the same read or read pairs, which is more likely to be caused by alignment errors; 5) candidates that are supported by more than 50% of the “high-quality” reads after confirming the alignment by a second aligner and masking bases adjacent to the start, end or spliced junctions of each read; 6) candidates that are recurrently present in the RNA-seq data of more than two unrelated individuals. The source code and default configuration file of RNA-MosaicHunter are publicly available at https://gitlab.aleelab.net/august/rna-mosaichunter.git, and it supports users to customize parameters that are used in the Bayesian genotyper and empirical error filters.

### Somatic calling from RNA-seq data

Each downloaded RNA-seq bam file was first converted back to the fastq format by Picard (v1.138) and then aligned to the GRCh37 human reference genome by STAR (v2.5.0a)^8^ in the two-pass mode, where the reference gene annotation (Gencode version 19) was used in the first pass and then a sample-specific annotation generated from the first pass was used in the second pass. The aligned reads were processed by Picard (v1.138) to remove duplicates, followed by SplitNCigarReads, indel realignment, and base quality recalibration of GATK (v3.6)^9^. Reads that were improperly paired or with ambiguous alignment were removed, and only genomic positions covered by 10 or more reads were subject to RNA-MosaicHunter. To exclude A-to-I(G) RNA editing sites, we only considered non-A-to-G candidates from the output of RNA-MosaicHunter. We further excluded non-exonic candidates and candidates that are present in the polymorphism databases of the general human population including dbSNP^10^, the 1000 Genomes Project^11^, the Exome Sequencing Project^12^, and the Exome Aggregation Consortium^13^.

### Benchmarking of RNA-MosaicHunter

RNA-seq and whole-exome sequencing data of 19 esophageal carcinoma samples as well as whole-exome sequencing data of their matched normal samples were downloaded from The Cancer Genome Atlas (TCGA) Research Network^14^. The list of 19 esophageal carcinoma samples is: TCGA-L5-A4OF-01A, TCGA-V5-A7RC-01B, TCGA-LN-A4A1-01A, TCGA-IG-A97I-01A, TCGA-L5-A8NE-01A, TCGA-JY-A93C-01A, TCGA-LN-A49M-01A, TCGA-IG-A3YB-01A, TCGA-LN-A49Y-01A, TCGA-L5-A8NN-01A, TCGA-LN-A49L-01A, TCGA-LN-A9FQ-01A, TCGA-L5-A4OR-01A, TCGA-LN-A8I1-01A, TCGA-L5-A891-01A, TCGA-L7-A6VZ-01A, TCGA-LN-A4A4-01A, TCGA-LN-A5U5-01A, TCGA-L5-A4OJ-01A.

Somatic mutation calls created by the Broad Institute through the comparison of tumor and matched normal whole-exome sequencing pairs using MuTect^15^ were also downloaded. A total of 851 non-A-to-G, autosomal, exonic, tumor-specific somatic mutations were called from the 19 tumor samples and covered by 10 or more reads in tumor RNA-seq data. This callset served as the gold standard for benchmarking our RNA-seq somatic mutation calling pipeline. We applied our calling pipeline to 19 esophageal tumor RNA-seq profiles, without applying a filter for removing recurrent candidates because these tumor samples may share common driver mutations, and identified 613 non-A-to-G somatic mutations.

By comparing the RNA-MosaicHunter callset with the gold standard, we found that RNA-MosaicHunter successfully identified 499 out of 851 MuTect-called mutations, equivalent to a sensitivity of 59% (Fig. 2b). On the other hand, among 613 RNA-MosaicHunter-called mutations, 513 were confirmed by the MuTect calls while 65 mutations were missed by MuTect but showed reads with 2% or more mutant allele fractions in the DNA-seq data, suggesting an overall precision of 94% for RNA-MosaicHunter (Fig. 2a-b).

### Neuronal proportion estimation

To estimate the proportion of neurons and other brain cell types in bulk brain RNA-seq data of ROSMAP and MayoRNAseq, we applied CIBERSORT (v1.05)^16^ to deconvolute the cell-type composition for each RNA-seq sample, by using the cell-type-specific expression reference for different neuronal and glial types (excitatory and inhibitory neuronal subtypes in the cortex, cerebellar granule cells, Purkinje cells, endothelial cells, pericytes, astrocytes, oligodendrocytes and their precursor cells, and microglia), generated from a large-scale brain single-cell RNA-seq dataset^17^. We summed the estimated proportion of all subtypes of excitatory and inhibitory neurons to calculate the overall neuronal proportion for each sample.

### Panel design and sequencing

For hybrid capture, probes targeting the exons and exon-intron junctions of 149 cancer driver genes (Supplementary Table 3) were designed using the SureSelect DNA Advanced Design Wizard. The list of targeted genes was designed to include frequently mutated oncogenes and tumor suppressor genes in various types of cancer and clonal hematopoiesis. A total of 23,171 probes with a genomic size of 691 kbp were eventually designed and generated. These probes were then used for gene capture followed by library preparation using the SureSelect XT HS2 DNA Reagent Kit with 30 ng gDNA input. All prepared libraries were sequenced using three Illumina NovaSeq 6000 S4 flow cells with 150 bp paired-end reads.

### Somatic mutation calling from panel sequencing

The UMI information of each read was first extracted from the fastq files by AGeNT’s Trimmer (v2.0.2), and then reads were aligned to the GRCh37 human reference genome by BWA-MEM (v0.7.15)^18^. The aligned reads were processed by AGeNT LocatIt (v2.0.2) to generate the consensus read sequence from multiple reads that were derived from the same original DNA fragment and thus carried the same UMI, followed by GATK’s indel realignment (v3.6)^9^. We only kept the consensus reads that were supported by two or more reads in both strands. As a result, we achieved comparable depth and coverage between the AD and control samples, with more than 1000X average depth and more than 80% coverage of the targeted regions at >500X for consensus reads (Supplementary Table 4 and Extended Data Fig. 2a).

sSNVs and somatic indels (sIndels) were called from the consensus reads by MosaicHunter (v1.0)^7^ and Pisces (v5.3)^19^, respectively. For sSNV, MosaicHunter calculated the likelihood of the presence of a mutant allele, and only the candidates with a 0.5 or higher likelihood, 100 or more total reads, and 4 or more mutant-supporting reads were considered. We further excluded candidates as germline mutations if i) they have a 30% or higher mutation allele fraction; 2) the counts of mutant-supporting and total reads do not significantly deviate from the binomial distribution for heterozygous mutations (*p* ≥ 0.05); 3) they are present in the polymorphism databases (dbSNP^10^, the 1000 Genomes Project^11^, the Exome Sequencing Project^12^, and the Exome Aggregation Consortium^13^) or have a 0.01% or higher population allele frequency in the Genome Aggregation Database^20^. sIndels were called by Pisces with its default parameters, and a similar method was used to call mutation candidates and remove germline mutations.

To balance the sensitivity and specificity of our sSNV and sIndel detection, we developed two different pipelines when considering the recurrent presence across multiple individuals. The “stringent” pipeline only kept the mutations that were detected in one sample and completely absent in any other samples, whereas the “sensitive” pipeline additionally allowed the mutations that were exclusively present or specifically enriched (two-sample Z-test of proportion with *p* < 0.05) in the AD or control group.

### Benchmarking of mutation calling using panel sequencing

A mixing experiment was performed to benchmark the performance of the designed panel and variant calling pipeline. Germline mutation calls from two unrelated individuals, NA12878 and NA24695, were downloaded from the website of the Genome in a Bottle Consortium^21^. Genomic sites in the covered regions of panel sequencing that were genotyped as heterozygous in NA24695 but reference-homozygous in NA12878 were considered as the gold-standard list of somatic mutations, and gDNA from these two individuals were mixed to reach 10%, 5%, 2%, 1%, 0.5%, and 0.2% mutant allele fractions for these mutations. We applied the same experiment and analysis protocols of panel sequencing to the mixed samples with varied allele fractions, and then checked the proportion of gold-standard mutations that were identified by our identification pipeline as well as the consistency between expected and observed allele fractions.

### Fluorescence-activated nuclei sorting (FANS)

Nuclei were prepared following the previously published work^22^. Briefly, fresh frozen human brain tissue samples were first lysed in a dounce homogenizer using a chilled nuclear lysis buffer (10mM Tris-HCl, 0.32M Sucrose, 3mM Mg(Acetate)2, 5mM CaCl2, 0.1mM EDTA, pH 8, 1mM DTT, 0.1% Triton X-100) on ice. Tissue lysates were layered on top of a sucrose cushion buffer (1.8 M sucrose 3 mM Mg(OAc)2, 10 mM Tris-HCl, 1 mM DTT, pH 8) and ultra-centrifuged for 1 h at 30,000g. Nuclear pellets we resuspended in ice-cold PBS supplemented with 3mM MgCl2, filtered, and then stained with the neuronal marker (NeuN, Millipore MAB377) or microglial marker (CSF1R, Cell Signaling 65396) together with DAPI. For each brain sample, neuronal (NeuN+), glial (NeuN-), microglial (CSF1R+), and total (DAPI+) nuclei populations were sorted into 96-well plates by flow cytometry.

### Cell type analysis from 10X snRNAseq

For the PFC sample of one AD patient (with a *TET2* p.Pro1194Ser sSNV) and one healthy control, ten thousand microglial nuclei were sorted separately into a well of the 96-well plate and used for droplet generation and sequencing library preparation using the 10X Genomics Next GEM Single Cell 3′ GEM Kit v3.1 and Chromium Controller, following the manufacturer’s manual. The snRNAseq libraries were sequenced by Illumina HiSeq X, and down-sampled to have a comparable sequencing throughput. We also downloaded a large-scale snRNAseq dataset^23^, consisting of 80,660 nuclei isolated from 24 AD and 24 control PFC samples collected by ROSMAP, to serve as the reference. The sequencing data of our AD and control sample was firstly processed by Cell Ranger (v6.0.0)^24^ and then integrated and analyzed along with the reference dataset by Seurat (v4.9.9)^25^, for variance normalization, anchor-based RPCA integration, PCA clustering, and UMAP visualization. Cell clusters were manually annotated into different cell types based on the expression profile of marker genes (Extended Data Fig. 4b) for the major brain^26^ and blood^27^ cell types (*HBA1*: red blood cell; *CD3E*: T-cell; *CCR7*: B-cell; *FCN1*: monocyte). Our snRNAseq result confirmed 75-77% microglia purity in the CSF1R+ sorted nuclei of the AD and control brains, with additional 4-9% CNS-associated macrophages (Fig. 4a). We also observed minimal blood contamination in the sorted microglial population, with only 1% monocytes and the absence of other major blood cell types including red blood cells, T-cells, and B-cells (Fig. 4a and Extended Data Fig. 4b). Using this reference dataset, we also confirmed the minimal contamination of blood cells (< 0.3%) in ROSMAP brain samples.

### Amplicon sequencing

Amplicon sequencing was performed for validation and mutant allele fraction estimation in both bulk gDNA samples and sorted nuclei. Bulk gDNA was extracted from frozen brain samples using the EZ1 DNA Tissue Kit (Qiagen 953034). Five hundred nuclei of each cell type from each brain sample were sorted into 96-well plates with four replicates. Whole-genome amplification was then performed for sorted nuclei using the ResolveDNA Whole Genome Amplification Kit (BioSkryb Genomics) to meet the minimal DNA amount for panel sequencing. For each identified sSNV, three sets of primers were designed for PCR amplification of the targeted genomic region. PCR amplification was performed using the Phusion Hot Start II DNA Polymerase kit (Thermo Fisher F549L) with the following cycles: 98 °C for 30 sec; 5 cycles of 98 °C for 10 sec, 68 °C for 30 sec (decrease 1 °C/cycle), and 72 °C for 30 sec; 25 cycles of 98 °C for 10 sec, 63 °C for 30 sec, 72 °C for 30 sec; 72 °C for 10 min. The annealing temperatures of primers varied for each design which was determined by a testing PCR. PCR products were then purified using AMPure XP beads (Beckman Coulter A63882) and pooled for Amplicon-EZ sequencing (GENEWIZ).

The sequencing reads were first aligned to the GRCh37 human reference genome by BWA-MEM (v0.7.15)^18^ and then processed by GATK (v3.6) for indel realignment^9^. For each somatic mutation candidate, the number of reads supporting each allele was calculated by MosaicHunter (v1.0) and manually verified by Integrative Genomics Viewer (v2.3.93)^28^. A candidate was considered validated as somatic mutation (Extended Data Fig. 2e-g) if 1) the read fraction of the mutant allele was more than three times as high as the fractions of the other two error alleles in all three amplicons (somatic-I); or 2) the read fraction of the mutant allele in the corresponding brain sample was significantly higher than the fraction in an unrelated negative control brain sample for all three amplicons (somatic-II).

### Functional annotation of sSNV and sIndel

ANNOVAR (v2015Mar22)^29^ was applied to annotate somatic mutations into different genic categories: 5’ UTR, exonic (coding sequence), 3’ UTR, splicing (within intronic 2 bp of a splicing junction), and intronic. Exonic somatic mutations were further classified into multiple categories based on their predicted impacts on amino acids. A somatic mutation was labeled as deleterious if 1) it was annotated as splicing or predicted to cause stop-codon gain/loss; 2) it was a frameshift insertion or deletion; or 3) it was a missense mutation whose amino acid change was predicted to be deleterious by either PolyPhen2^30^ or SIFT^31^. For 149 cancer driver genes, we grouped them into (proto-)oncogenes and tumor suppressor genes (TSGs) according to the annotation of the COSMIC Cancer Gene Census^32^. Genes annotated as both oncogenes and TSGs were not considered in calculating the mutation burdens plotted in Fig. 3d. MAFTools (v2.10.1)^33^ was used to illustrate the gene-level distribution of somatic mutations. Genes and driver mutations involved in clonal hematopoiesis of indeterminate potential (CHIP) were extracted from a study that analyzed blood whole-genome sequencing data from 11,262 people^34^.

Functional enrichment analysis of Gene Ontology (GO) terms was performed using GOseq (v1.34.1)^35^. Exonic somatic mutations identified from the RNA-seq of AD patients or normal controls were used as the input, and Wallenius’ noncentral hypergeometric distribution was used to test the enrichment, with a probability weighting function to control for potential gene length bias. Only GO terms with 3 or more hits and an initial overrepresentation p-value < 0.01 were considered. GO terms with more than 1000 genes were excluded. All the GO terms with significant enrichment of AD somatic mutations were plotted in Fig. 2f, where the p-value was adjusted by Hommel’s method for the correction of multiple hypothesis testing. In comparison, only one GO term “helicase activity” showed significant enrichment for somatic mutations identified from normal controls.

### Burden analysis of sSNV and sIndel

Somatic mutation density in each clinical group was calculated by counting the total number of somatic mutations and dividing it by the total size of powered genomic regions with ≥10X coverage for RNA-seq or ≥500X for panel sequencing data sets, and the odds ratio and the two-sample Z-test of proportion were used to test whether the AD group had a higher mutation burden than the control group. In the gene-level analysis for panel sequencing data, we compared the somatic mutation burden between AD and control groups using a similar two-sample Z-test of proportion, in which the total genomic size for each gene was calculated as the product of the exonic length and the number of individuals in AD or control group.

For the linear regression analysis, the count of somatic mutations in each sample was modeled as a continuous outcome, whereas clinical status and other covariates of interest (e.g. age, sex, sequencing depth, post-mortem interval, and neuronal proportion) were modeled as independent variables. Our linear regression results from both RNA-seq and panel sequencing confirmed the increased burden of somatic mutation in AD brains after controlling for all of these potential confounding factors (Fig. 2e and 3c). We only considered donors with ages less than 90, because all the donors with age 90 or higher were labeled as “90+” in the demographic tables of the ROSMAP and MayoRNAseq studies. We also tested whether APOE4 carriers exhibited different somatic mutation burdens compared to non-carriers by considering this as an additional covariate. However, the known strong correlation between the APOE4 allele and AD risk may violate the independence of covariate assumption in linear regression, thus limiting the statistical power. To further rule out the effect of potential blood contamination, we measured the normalized gene expression level (transcript per million, TPM) of blood marker genes including *HBA1*, *CD3E*, *CCR7*, and *FCN1* for each RNA-seq sample of ROSMAP and MayoRNAseq by StringTie (v1.3.3b)^36^, and then modeled them as additional covariates in our linear regression model. We observed minimal contamination of blood-derived immune cells in ROSMAP and MayoRNAseq brain samples, and confirmed that our observed AD increase remains significant after controlling for any of these genes (*p* ≤ 0.01).

### Positive selection analysis

Signals of positive selection were assessed for sSNVs identified from AD and control samples separately by dNdScv^37^. The dN/dS ratios and p-value for missense, nonsense, and splicing mutations were calculated at the levels of individual genes and groups of genes, by comparing against the background synonymous mutation rate with the consideration of the sequence composition of genes. For each gene in AD or control group, we 1) calculated the number of missense and truncating (nonsense and splicing) mutations under positive selection by multiplying the number of all mutations in that gene by the proportion of positively selected mutations inferred from the gene-specific dN/dS ratio; 2) determined the proportion of positively selected cells by multiplying the number of positively selected mutations by the average mutant allele fraction in that gene × 2 (given that almost all the sSNVs should be heterozygous in carrier cells). Assuming a consistent number of profiled cells in panel sequencing for each brain, we further estimated the number of positively selected cells in each AD and control brain by aggregating the number of positively selected cells across the group of genes and normalizing this number based on the count of brain samples in AD and control groups.

### Automatic cell-type identification with scType

Myeloid cells in the brain include both parenchymal microglia and CNS-associated macrophages (CAMs), including meningeal, choroid plexus, and perivascular macrophages (PVMs)^38^. Microglia-perivascular macrophages, hereby referred to as microglia-CAMs, represented 3.37% of all pre-annotated cells within SEA-AD, which is slightly lower than past estimates of microglia-CAMs making up 5-15% of all brain cells^39,40^. scType (v20220909)^41^ was used to automatically identify any additional high-quality microglia-CAMs beyond those originally annotated in SEA-AD (“pre-annotated” cells) to increase statistical power for calling mosaic chromosomal alterations (mCAs). Excitatory neurons (ExNs) were also automatically typed as a cell-type out-group to further facilitate accurate identification of microglia-CAMs, as scType’d microglia-CAMs should have high microglia-CAM scType scores but low ExN scType scores.

Prior to running scType, each SEA-AD sample was processed, normalized, and clustered with the Louvain algorithm using Seurat (v4.1.1)^25^. Each sample underwent quality control with the following metrics: retain only 1) genes expressed in ≥ 3 cells, 2) cells with ≥ 10 expressed genes, 3) cells with ≤ 5% mitochondrial gene expression, 4) cells with > 250 expressed genes and < 7500 expressed genes. Positive markers for microglia-CAMs (*P2RY12, ITGAM, CD40, PTPRC, CD68, AIF1, CX3CR1, TMEM119, ADGRE1, C1QA, NOS2, TNF, ISYNA1, CCL4, ADORA3, ADRB2, BHLHE41, BIN1, KLF2, NAV3, RHOB, SALL1, SIGLEC8, SLC1A3, SPRY1, TAL1*) and ExNs (*SLC17A7, SLC17A6, GRIN1, GRIN2B, GLS, GLUL, GRIN2A*) were downloaded from the scType marker database and used to calculate microglia-CAM and ExN scType scores for each individual cell.

In brief, scType calculates cell-type specific scores for each cell using a weighted and normalized aggregation of marker gene expression, where marker genes are weighted more highly if they are more specific for a given cell type (expressed in one cell type of interest, rather than several). For each sample, both ExN and microglia-CAM scType scores were calculated for cells that were pre-annotated as either ExNs or microglia-CAMs. Taking these pre-annotations as ground truth, ROCR (v1.0.11)^42^ and cutpointr (v1.1.2)^43^ were used to calculate the optimal cutpoint for ExN and microglia-CAM scType scores that maximized the sum of sensitivity and specificity of classification over 1000 bootstraps. Using these learned ExN and microglia-CAM cutpoints, cells that were not pre-annotated were assigned as ExNs, microglia-CAMs, or neither. A small number of cells had both microglia-CAM and ExN scType scores greater than the corresponding optimal cutpoints; these cells were discarded due to ambiguity in assignment.

In addition to filtering of individual cells, 6 samples were filtered out due to not meeting at least one of the following sample-specific metrics: 1) microglia-CAM AUC > 0.9, 2) ExN AUC > 0.9, 3) fraction of pre-annotated ExN typed by scType as microglia < 0.1, and 4) total number of pre-annotated and scType’d microglia-CAMs > 50. This analysis filtered one individual *H20.33.008*, as this donor had only one associated sample that was filtered due to not meeting the above sample-specific metrics.

As a final step to ensure that scType’d cell microglia-CAMs were highly similar to their corresponding pre-annotated cell types, pre-annotated and scType’d microglia-CAMs derived from the same donor were merged into a single Seurat object and processed, normalized, and clustered using the Louvain algorithm. Clusters in which over 50% of cells were pre-annotated microglia-CAMs were identified and only scType’d microglia-CAMs in these clusters were retained as high-confidence scType’d microglia-CAMs cells. Only pre-annotated microglia-CAMs and these high-confidence scType’d microglia-CAM cells were used for mCA-calling and all subsequent downstream analyses.

### mCA calling from snRNAseq

Genomic regions of non-uniparental disomy CH-associated mCA listed in Extended Data Figure 4d and 4e of Saiki *et al.*^44^ were extracted, and genomic coordinates of these regions were downloaded from the hg38 reference genome accessed through the UCSC Genome Browser^45^.

mCA calling was done for microglia-CAM, astrocytes, oligodendrocytes, oligodendrocyte precursor cells (OPCs), and ExNs. For each cell type, raw count matrices (gene × cell) were extracted for the 31 AD cases and 31 age-matched healthy controls that passed filtering as described above. Each of these matrices was processed and normalized using Seurat (v4.1.1) and then independently used as input for mCA-calling with CONICSmat (v0.0.0.1)^46^.

The aforementioned mCA regions identified in Saiki et al., were tested with CONICSmat (Supplementary Table 7), and raw mCA calls were further filtered to increase specificity of calls. In brief, a putative mCA was retained if it met the following criteria: 1) Bonferonni adjusted p-value < 0.05; 2) <25% ambiguous cells (cells with a posterior probability > 0.25 and < 0.75); 3) median expression of putative mCA-carrying cells is > or < 1.96 standard deviations of putative normal cells of the same type for amplifications or deletions, respectively; 4) no negative control regions (i.e. whole chromosome regions that have not been associated with mCA in past literature) showed a larger difference in expression between putative normal and mCA-carrying cells than the called mCA; 5) the expression of putative normal cells was within 1.96 standard deviations of baseline expression of cells of the same type across all other individuals; and 6) the same mCA was not called in a different cell-type from the same individual. For microglia-CAMs, putative mCAs were additionally filtered if the number of scType’d non-ambiguous cells (posterior probability < 0.25 or > 0.75) were ≤ 1.5x the number of pre-annotated non-ambiguous cells for both altered and wild-type cells. This filtering criterion was added to ensure that mCA calls identified from scType’d and pre-annotated microglia-CAMs were not driven by added scType’d cells.

### Burden analysis of mCA

Per cell type, the number of cells with mCAs from AD donors, the number of cells without mCAs from AD donors, the number of cells with mCAs from control donors, and the number of cells without mCAs from control donors were counted and an odds ratio (OR) of mCA-carrying cells in AD donors vs control donors was calculated. For two cell types, CAMs and oligodendrocytes, all mCA-carrying cells were in AD donors and the OR was thus infinite. To facilitate comparison of the actual OR against an empirical null as described below, a pseudocount of 1 was added to the number of mCA-carrying cells in AD and control groups separately for these two cell types. To calculate the significance level of this calculated odds ratio, an empirical null was generated using permutation. In brief, for each cell type, diagnosis labels were permuted over the set of all cells from each donor, including both mCA-carrying and wild-type cells. If a donor had multiple called mCAs, diagnosis labels were permuted over each mCA individually. Specifically, for each called mCA in a given individual, cells were divided into wild-type or mCA-carrying for that specific mCA. Each of these partitions of wild-type versus mCA-carrying cells was then randomly assigned a diagnosis status. OR was calculated for each set of permutated data. Permutations were repeated 1000 times and the p-value of the actual OR was calculated as 1 – the percentile rank of actual OR against the empirical null distribution of permutation ORs. Ten trials of 1000 permutations were completed to ensure the robustness of p-values.

### Creation of an integrated snRNAseq microglia-CAM atlas

All scType’d and pre-annotated microglia-CAMs from AD and healthy control samples, with the exception of the one associated with *H20.33.008* as described above, were individually processed with Seurat (v4.1.1). In brief, each sample underwent quality control with the following metrics: retain only 1) genes expressed in ≥ 3 cells, 2) cells with ≥ 10 expressed genes, 3) cells with ≤ 5% mitochondrial gene expression, 4) cells with > 250 expressed genes and < 7500 expressed genes. Variance-stabilizing normalization and regression of the technical covariates percent.mt, nFeature_RNA, and nCount_RNA were performed with Seurat function SCTransform, and clustering was done using the Louvain algorithm.

Individual samples were then merged into a single Seurat object, and dimensionality reduction was performed using PCA. This merged object was then integrated over constituent individual samples using Seurat’s wrapper function for Harmony (v0.1.1)^47^. UMAP visualization of the integrated object showed no visible clustering by sample ID or individual ID, consistent with successful integration (Extended Data Fig. 6).

### Differential expression analysis and functional annotation of integrated microglia-CAM snRNAseq atlas

Differential expression analysis was performed between microglia-CAMs with and without called mCAs from mCA-carrying AD individuals using the FindMarkers function of Seurat (v4.1.1) with a min.pct cutoff of 0.10 and no fold-change cutoff. Genes with an adjusted p-value < 0.05 were called as differentially-expressed genes (DEGs).

clusterProfiler (v4.4.4)^48^ was used to perform all enrichment analyses. GO enrichment analysis was performed using standard parameters and a universe of all genes expressed in >10% of microglia-CAMs in the integrated atlas. Terms were deemed significant if they had an adjusted p-value < 0.05.

DEGs were also tested for enrichment of previously defined microglial state gene modules^49^. A minority of genes (107/905; 11.9%) within these microglial state gene modules were shared between multiple modules. To ensure specificity of module enrichment, genes were weighted by the inverse of the number of modules in which they were present. Non-integer values were rounded and module enrichment was tested using a hypergeometric test.

### Data and material availability

All the RNA-seq and DNA-seq data of ROSMAP, MayoRNAseq, and SEA-AD are available via the AMP-AD Knowledge Portal. The RNA-seq and DNA-seq data of TCGA are available via the NCI Genomic Data Commons Data Portal. ROSMAP resources can be requested at https://www.radc.rush.edu. The panel sequencing and snRNAseq data generated in this study will be deposited to the AMP-AD Knowledge Portal, with controlled use conditions set by human privacy regulations. Other materials are available from the authors upon reasonable request.

### Code availability

The source code and default configuration file of RNA-MosaicHunter are available at https://gitlab.aleelab.net/august/rna-mosaichunter.git. Custom bash and R scripts used in this study will be publicly available at https://gitlab.aleelab.net/august/ad-clonal.git.

## Supplementary Discussion

In this study, we observed that AD brain samples harbor an increased burden of somatic mutations in cancer driver genes, especially in CH-associated genes, suggesting that CH mutations in the brain are positively associated with AD pathogenesis. However, a study by Bouzid *et al.*^1^ finds that CH mutations in blood appear to be protective against AD. Another work from Kessler *et al.*^2^ reports no association between CH mutations in blood and AD risk in a much larger number of samples. Several technical and methodological differences may explain the inconsistency between these three studies.

First, our study was designed to directly study brain samples of AD patients and healthy controls, whereas both Bouzid *et al.* and Kessler *et al.* were based on the re-analysis of peripheral blood sequencing data. Although both studies reported that many of these CH mutations were shared between brain (microglia) and blood samples of the same individuals, it remained unclear whether CH mutations might have a different role in AD between the brain and blood (harmful in brain vs. protective/neutral in blood).

Second, we screened for brain somatic mutations by ultra-deep panel sequencing with a UMI design, such that we were able to detect mutations with MAFs as low as 0.1% (Extended Data Fig. 2). In comparison, Bouzid *et al.* and Kessler *et al.* utilized existing blood whole-exome sequencing data with conventional depth, which was designed for germline variant detection and could only detect CH mutations with MAFs > 5-10%^2,3^, although CH mutations with lower MAFs are more typical in the blood^4^. Indeed, we observed that the AD enrichment of somatic mutations in CH-associated genes disappears when only high-MAF mutations are considered (Extended Data Fig. 7b).

Finally, our panel sequencing covered a comprehensive list of 149 cancer driver genes (Supplementary Table 3), including many genes that had been reported in cancer development but not yet linked to CH. Our results suggest that somatic mutations in these non-CH-associated genes also show an increased burden in AD brains, robust with different MAF cutoffs (Extended Data Fig. 7a), but such effects would be missed in Bouzid *et al.* and Kessler *et al.* because their studies only focus on CH-associated genes.

